# Stage-Specific Modulation of Multinucleation, Fusion and Resorption by the Long Non-coding RNA DLEU1 and miR-16 in Human Primary Osteoclasts

**DOI:** 10.1101/2023.10.24.563436

**Authors:** Sara Reis Moura, Ana Beatriz Sousa, Jacob Bastholm Olesen, Mário Adolfo Barbosa, Kent Søe, Maria Inês Almeida

## Abstract

Osteoclasts are multinucleated cells formed through fusion of mononucleated precursors of the myeloid lineage and are the only cells that can resorb all the constituents of the bone matrix. Our goal was to investigate the role of long non-coding RNA *DLEU1* and miR-16-5p in the fusion of human primary osteoclasts and their resorptive capacity. We found *DLEU1* to be markedly upregulated, whereas miR-16 was significantly suppressed, during osteoclast differentiation, suggesting a potential involvement in the multinucleation process. The knockdown of *DLEU1* or the overexpression of miR-16 in human primary pre-osteoclasts from male human donors (50 years or older) impaired fusion at both early and late time-points, each in distinct ways, without affecting cell viability. Time-lapse recordings confirmed the impairment of the fusion process and showed an abrogation of the phagocytic cup fusion modality, as well as a reduction of the fusion between mononucleated precursors and multinucleated osteoclasts during *DLEU1* silencing. Furthermore, mass spectrometry-based quantitative proteomics revealed that the effects of *DLEU1* and miR-16 on osteoclast fusion were mediated by distinct proteins and processes. Thus, both *DLEU1* inhibition and/or miR-16 overexpression hinder osteoclast fusion through modulation of different mechanisms. Moreover, decreased levels of *DLEU1* specifically affected the resorption speed of pit-making osteoclasts, while increased levels of miR-16 promoted bone resorption mainly through pit-formation, impairing the resorption speed of the osteoclasts making trenches and affecting their resorbed area. Taken together, these findings identify *DLEU1* and miR-16 as mediators of osteoclast fusion and activity, offering potential new therapeutic targets to ameliorate bone destruction in skeletal diseases with accentuated bone deterioration.

## Introduction

Osteoporosis is a consequence of skeletal deterioration, characterized by alterations in the tissue microarchitecture and quality loss, resulting in increased vulnerability to fractures [1]. This condition is more prevalent in individuals over 50 years of age and is caused by a disruption in the delicate balance between bone formation and resorption [2,3]. In osteoporotic patients, the resorptive activity of the osteoclasts (OCs) surpasses the bone formation capacity, being a hallmark of this disease [4].

OC formation comprises several stages, including precursor recruitment, differentiation, fusion and activation / maturation. Each stage is essential and co-dependent for the proper functioning of the subsequent steps and ultimately, determine the OC’s resorptive capacity [5,6]. Therefore, any impairment in the early steps will have a subsequent influence on the later stages. In fact, a positive correlation between the nucleation status and the OC’s bone resorptive activity has already been established *in vitro* [7–9]. Interestingly, this association between hyperactivation of multinucleation and increased bone resorption by the OC is also observed in patients with Paget’s disease, leading to exacerbated bone resorption [10–15]. In addition, a recent study reported that the fusion potential of human OCs *in vitro* was found to be correlated with age, menopause status and bone resorption levels *in vivo* [16]. These findings reinforce the fact that the relationship between the fusion capacity and the resorption levels is not only observed *in vitro*, but also *in vivo*. Consequently, the regulation of bone destruction can be achieved either through modulation of preceding stages or directly at the resorptive stage. Mature OCs can remove bone tissue through two distinct strategies with very contrasting characteristics [17,18]. On one hand, there is the pit mode, in which an intermittent, restricted and less aggressive erosion is observed [17,18]. On the other hand, the trench mode is characterized by more aggressive bone resorption due to longer periods of bone resorption, faster erosion rates, deeper cavities and sensitivity to CTSK inhibition [17,18]. In fact, inhibition of CTSK has been described to abolish trench formation, while enhancing pit formation [19,20]. Considering the different degrees of aggressiveness associated with each resorption mode and their association to bone resorption *in vivo* [7], it seems appropriate to assume their potential clinical relevance. Therefore, besides evaluating the overall bone resorption activity, it is also of interest to investigate the impact of non-coding RNAs (ncRNAs) on both resorption modes separately.

During the last decades, several factors, both intrinsic and extrinsic, have been identified as key players in the cascade of events that culminate in OC maturation, activation and activity. Among them are NFATc1 (Nuclear Factor Of Activated T Cells 1) [21–23], DC-STAMP (dendritic cell-specific transmembrane protein) [24–29], CD47 [30–35], syncytin-1 [30,36] and e-cadherin [37,38]. Recently, ncRNAs have received more interest due to their abnormal expression profiles [39], often associated with disorders (*eg*. bone disorders) [40] and their ability to modulate distinct cellular processes involved in bone remodeling [2,41–45]. Nevertheless, a deep and extensive understanding of the role of specific microRNAs (miRNAs) and more importantly, long non-coding RNAs (lncRNAs), is needed for future clinical translation.

In this study, we investigate the role of small and long ncRNAs located in the chr 13q14.2-q14.3 region, which encodes *DLEU1* (Deleted In Lymphocytic Leukemia 1), *DLEU2* (Deleted In Lymphocytic Leukemia 2), miR-15 and miR-16, during the distinct stages of OCs maturation, including fusion and resorption [46]. Currently, the role of these closely located ncRNAs in non-tumorigenic-derived bone diseases and/or cells, specifically on OCs, has not yet been disclosed. Here, we perform an in-depth analysis on the OC phenotypes regulated by *DLEU1* and miR-16, in a stage-specific manner. Altogether, our findings reveal an antagonistic relationship between *DLEU1* and miR-16, when considering OC fusion. We also provide comprehensive insights into the impact of these molecules on individual fusion events. Moreover, our study highlights distinct effects for *DLEU1* and miR-16 on bone resorption, with *DLEU1* primarily influencing the resorption speed during pit-formation, while miR-16 is shown to affect both resorption modalities.

## Materials and Methods

### *In vitro* generation of human OCs from buffy coats

For osteoclastogenic differentiation and functional assays, CD14^+^ monocytes were purified, as previously reported, from human buffy coats (BC) of anonymous 50-65 years old male blood donors, with their informed consent, donated by the Odense University Hospital (Denmark) [7,47,48]. Briefly, BC were diluted, laid over Ficoll-Paque (Amersham, GE Healthcare, Little Chalfont, UK) and centrifuged at 900 *g*, for 20 min. Peripheral Blood Mononuclear Cells (PBMCs) were then collected, washed and CD14^+^ monocytes were subsequently isolated by immunomagnetic separation, using the BD IMag Anti-Human CD14 Magnetic Particles – DM (BD Biosciences, San Jose, CA, USA), according to the instructions given by the supplier. Cells were seeded at a cell density of 6.7 × 10^4^ cells/cm^2^ in cell culture flasks (Greiner), in α-MEM (Thermo Fisher Scientific), supplemented with 10 % (v/v) FBS (Sigma-Aldrich), 1 % (v/v) Penicillin-Streptomycin (P/S, Sigma-Aldrich) and 25 ng/mL M-CSF (R&D Systems) for 2 days. Afterwards, the culture media was renewed and cells were differentiated into mature osteoclasts (OCs) over the course of 7 additional days with 25 ng/mL M-CSF and 25 ng/mL RANKL (both from R&D Systems), as previously described [17,30,49]. An independent cohort of BC from healthy donors was collected at Hospital Universitário São João (Portugal) for validation experiments, and isolation and cell culture of these donors was performed as already reported [41].

### Oligonucleotides transfection

Human primary OCs in several stages of differentiation were transiently transfected with either a silencing RNA against Deleted In Lymphocytic Leukemia 1 (DLEU1) (siDLEU1; 25 nM; Lincode Human DLEU1 siRNA – SMARTpool; R-0200009-00-0005; Dharmacon), miR-16 mimic (miR-16; 50 nM; mirVana® miRNA mimic; MC10339; ThermoFisher Scientific), or the respective negative controls [Lincode Non-targeting Pool (CTR lncRNA; 25 nM; D-001320-10-05; Dharmacon) and mirVana™ miRNA Mimic, Negative Control #1 (CTR mimics; 50 nM; 4464058; ThermoFisher Scientific)].

The oligonucleotides were delivered using the GenMute™ siRNA transfection reagent for primary macrophages (SL100568-PMG, SignaGen Laboratories), according to the manufacturer’s protocol. Briefly, a working solution was prepared, the oligonucleotides were individually added, according to the concentrations mentioned above, and the solution was left to incubate for 15 min, to allow the transfection complexes to form, and then added to the cells.

The transfections were performed at day 3 post-monocyte isolation for fusion assays, RNA and protein collection; at day 4 post-monocyte isolation for fusion time-lapses recordings; and at day 8 post-monocyte isolation for resorption experiments (both end-point and time-lapse recordings).

### RNA isolation

The RNA was extracted from human primary OCs using the TRIzol Reagent (Invitrogen), according to the manufacturer’s instructions. RNA concentration and purity were determined using NanoDrop Spectrophotometer ND-1000 (ThermoFisher Scientific). Total RNA samples were stored at -80 °C until further use.

### Reverse transcription and real-time quantitative polymerase chain reaction (RT-qPCR)

For the analysis of the miRNA expression levels, TaqMan MicroRNA Reverse Transcription Kit and gene specific stem-loop Reverse Transcription primers (Applied Biosystems) were used, according to the manufacturer’s protocol and as previously described [41,42]. I-miR-15a-5p and I-miR-16-5p FASTA format sequence annotations were obtained from the miRbase database (http://www.mirbase.org/, Supplemental Table I). The RT-qPCR reaction was performed under the following conditions: 10 min at 95 °C, 40 cycles of 15 s at 95 °C and 1 min at 60 °C. Small nuclear RNA U6 was used as a reference gene.

For the analysis of the expression of coding and non-coding transcripts, RNA samples were digested with TURBO DNA-free kit (Life Technologies), following the manufacturer’s instructions, to remove potential DNA contaminants. Buffer and Dnase were added to the samples and incubated for 30 min, at 37 °C. Then, Dnase inhibitor was added to the samples, which were subsequently vortexed and centrifuged at 10 000 *g*, for 1 min. The supernatant containing the RNA was collected and the RNA samples were then used to synthesize cDNA, using random hexamers (Life Technologies) and the SuperScript® III Reverse Transcriptase kit (Life Technologies), as already described [41,42]. qPCR reaction was performed by adding SYBR Green PCR Mastermix (Bio-Rad), nuclease free water and the pair of primers corresponding to the transcript of interest (reverse and forward; Supplemental Table II), to the cDNA, as reported [50]. The reaction was performed in a CFX Real-Time PCR Detection System (Bio-Rad), according to the following conditions: 3 min at 95 °C (denaturation) and 40 cycles of 30 s at 95 °C (cDNA denaturation), 30 s at 58 °C (annealing) and 30 s at 72 °C (extension). Primer sequences are described in Supplemental Table II). B-ACTIN was used as a reference gene.

Relative expression levels for each targeted miRNA and genes were calculated using the quantification cycle (Cq) method, according to the MIQE guidelines [51]. Data were analyzed using the Bio-Rad CFX Manager software and all the qPCR reactions were performed in duplicate.

### Fusion assay

At day 5 and 9 of the differentiation process, OCs were fixed with 3.7 % (v/v) formalin solution and stained with Giemsa and May-Grünwald, as previously described [52]. After fixation, methanol was added to each well and left to incubate for 15 min. Cells were air dried and the May-Grünwald solution (J.T.Baker^®^), followed by the Giemsa solution (Merck), was added to the samples. Finally, cells were washed with distilled water twice and all the multinucleated OCs (≥2 nuclei), and their respective number of nuclei, were systematically counted in 7 different random fields (assigned by a random number generator) in 8-10 replicate wells for each condition, using the Axiovert 200 microscope (Carl Zeiss).

### Bone resorption assays and quantification of bone resorption

Mature OCs (cultured for 2 days with 25 ng/mL M-CSF and for 7 additional days with 25 ng/mL M-CSF and 25 ng/mL RANKL) were detached using accutase (Biowest). Subsequently, they were reseeded on top of 0.4 mm thick bovine cortical bone slices (Boneslices.com) to offer a biologically relevant bone substrate, surpassing the performance of synthetic materials or dentine slice. The seeding density was 5.0 x 10^4^ cells/bone slice (5 bone slices/condition). OCs were then cultured in α-MEM with 10 % (v/v) FBS, 25 ng/mL M-CSF and 25 ng/mL RANKL for 3 additional days [47]. Before the analysis of the eroded surface and cell lysis / removal, the cellTiter-Blue® Cell viability Assay was performed. Resorption events were visualized and quantified after staining with toluidine blue, as previously described [53]. Samples were analyzed by light microscopy for the percentage of eroded surface, type of resorption pattern (pits or trenches, in accordance with published definitions [17,49]) and the number of resorption events, with a 10x objective through a 100 point grid graticule (catalog number:01A24.5075, Graticules Optics) [18,49]. All quantifications were performed blinded with respect to the treatment.

### CellTiter-blue assay

To measure cells’ viability and activity, the CellTiter-Blue® Cell viability Assay (Promega) was used. Briefly, 10 % (v/v) of the dye was added to each well and left to incubate for 20 min, at 37 °C and 5 % (v/v) CO_2_, protected from the light. The supernatant was then collected, transferred to a black 96-well plate (Greiner) and the fluorescence was read at an excitation wavelength of 530 nm and an emission wavelength of 590 nm, using the spectrophotometer microplate reader Synergy MX (Biotek Synergy).

### Colorimetric lactate dehydrogenase (LDH) assay

Cell viability / cytotoxicity was also evaluated through quantification of the plasma membrane damage using the The CytoTox 96® Non-Radioactive Cytotoxicity Assay (Promega). The protocol was performed according to the manufacturer’s instructions. Briefly, the conversion of iodonitrotetrazolium violet (INT) into a red formazan product by LDH was measured through incubation with the substrate solution for 30 min, at RT, protected from the light. Then, a stop solution was added, and absorbance was measured at 490 nm, using the spectrophotometer microplate reader Synergy MX (Biotek Synergy). LDH levels in the conditioned-media were measured at day 5 of the differentiation process from OCs transfected at day 3.

### Protein extraction and quantification

Pre-OCs transfected with siDLEU1, miR-16 mimics, or the respective controls, at day 3 were cultured under osteoclastogenic differentiation conditions for 2 additional days. Cells were lysed in the presence of phosphatase and protease inhibitors (Thermo Scientific) at day 5 of the differentiation process, after being washed with PBS 1x. Cell lysates were clarified through a 20 000 *g* centrifugation, for 10 min at 4°C. The supernatants, containing the protein, were collected and protein concentration was determined using DC Protein assay kit (Bio-Rad).

### Proteomic analysis

For the mass-spectrometry analysis, 40 μg of protein lysates from transfected OCs were used. Protein identification and label-free quantitation were performed by nanoLC-MS/MS, composed by an Ultimate 3000 liquid chromatography system coupled to a Q-Exactive Hybrid Quadrupole-Orbitrap mass spectrometer (Thermo Scientific), as previously reported [41,50,54]. Raw data were processed using Proteome Discoverer 2.3.0.523 software (Thermo Scientific). Protein identification was performed with Sequest HT search engine against the *Homo sapiens* entries from the UniProt database (https://www.uniprot.org/). Protein and peptide confidence were set to high. Analyzed samples were normalized against the total peptide signal and its quantitative evaluation was achieved by pairwise comparisons of the detected peptides. Data were corrected using the Benjamin Hochberg method. Protein levels were compared using the median ratio.

A minimum of two unique peptides per identified protein were required for further evaluation and contaminants were removed. Bioinformatics and data analysis were performed with a *p*<0.05. The differentially expressed proteins were classified according to the Gene Ontology (GO) annotations and enriched pathways. The study of the terms enriched in our differentially expressed proteins was performed using Protein Analysis Through Evolutionary Relationships (PANTHER) tool [55–60]. Gene Set Enrichment Analysis (GSEA) was also used for interpreting the expression data [61,62].

### Time-lapse recordings and analysis of fusion assays

Three days after being reseeded into eight-wells of a Nunc Lab-Tek II chambered cover-glass (Nunc−Thermo Fisher Scientific) at a density of 6.0x10^4^ cells/well in α-MEM, supplemented with 10 % (v/v) FBS, 25 ng/ml M-CSF and 25 ng/ml RANKL, and one day after being transfected with either siDLEU1, miR-16 mimics, or the respective controls, the OCs reached an early fusion stage (confirmed through light-microscopy) and the time-lapse recordings were initiated (day 5). The chambered cover-glass was subsequently placed in the incubation chamber of the confocal Olympus Fluoview FV10i microscope (Olympus Corporation) with 5 % (v/v) CO_2_ at 37 °C, for 4 days (between days 5 and 9 of the differentiation process). The media was changed 2 days after the beginning of the recordings (day 7). For each condition, four random sites were recorded (in a total of 16 different sites over the course of 4 days). The time-lapse images were acquired every 21 min for about 24 h, using phase contrast. This procedure was repeated for four consecutive days and for each new recording four new sites were chosen in each well.

Analysis of the time-lapse recordings for the measurement of migration and cell’s shape/size, as well as to observe any fusion events, was performed using the FV10-ASW 4.1/4.2 Viewer software (Olympus) and ImageJ [63]. When a fusion event was detected, it was characterized as previously reported [32,64].

Data was collected from four independent donors, all with the four different experimental conditions (siDLEU1, miR-16 mimics, or the respective controls). For each donor, a total of 64 videos, reflecting a total of 1 536 hours, were analyzed.

### Time-lapse recordings and analysis of resorption assays

At day 8 of the differentiation process, mature OCs were transfected with either siDLEU1, miR-16 mimics, or the respective controls. On the next day, the OCs were detached, labelled with 100 nM SiR-actin and 10 μM verapamil (both from Spirochrome), as already described [17] and reseeded onto 0.2 mm bone slices labelled with N-hydroxysuccinimide ester-activated rhodamine fluorescent dye (ThermoFisher Scientific), at a cell density of 1x10^5^ cells/bone slice (Boneslices.com), in 96-well plates [17,65]. Afterwards, the bone slices and the labelled transfected mature OCs were placed into eight-wells of a Nunc Lab-Tek II chambered cover-glass and bone resorption events were recorded using the confocal Olympus Fluoview FV10i microscope, at 37 °C, in 5 % (v/v) CO_2_, in a humidified atmosphere. OCs were imaged using a 10x objective and a confocal aperture of 2.0. For each condition and donor, 3 random sites were recorded for 3 uninterrupted days, according to previous publication [17].

Analysis of the time-lapse recordings for the measurement of the resorbed area and time, characterization of the resorption events and cell’s size, was performed using the FV10-ASW 4.1/4.2 Viewer software (Olympus) and ImageJ [63]. The duration of each event was determined by calculating the number of frames elapsed from the start to the conclusion of that specific event. When resorbing in pit-mode, OCs display a round actin ring, which remains stationary during the resorption process. These resorption events were identified by the initial presence of an F-actin staining at the center of the actin ring, indicating the formation of the ruffled border, along with the start of labelled collagen removal from the bone slice surface. The termination of these resorptions events was determined by either the cessation of the ruffled border, the disappearance of the actin ring or the relocation of the OC from the event. In case of the OC resorbing in trench mode, it is observed an initial displacement of the actin ring towards a specific side of the pit (if the trench formation started after pit generation) or initiation of collagen removal that is expanding only in one direction (when the OC initiate the resorption process in trench mode right from the start). Trenches that started outside the recording window were solely used for calculating the overall resorption speeds and were not categorized based on their initiation, due to the unavailability of this data. The resorbed area of each event was determined by manually outlining the boundaries of the resulting resorption cavities, represented by the ‘black’ regions on the Rhodamine staining, using ImageJ.

Data was collected from three different donors, all with the four different experimental conditions (siDLEU1, miR-16 mimics or the respective controls). For each condition and donor, a total of 216 h were analyzed (648 h when considering the four conditions).

### Statistical analysis

All graphs were performed using GraphPad Prism software, version 9 (GraphPad software). Firstly, data sets were tested, using the Shapiro-Wilk and Kolmogorov-Smirnov normality tests to determine if the data followed a parametric or non-parametric distribution. When the data passed the normality tests, a student *t* test (2 groups) or one-way ANOVA (> 2 groups), followed by Sidak’s multiple comparison or Turkey’s multiple comparisons tests were used. For non-normal distribution data, non-parametric tests were used to evaluate significant differences between samples, namely two-tailed Wilcoxon matched pairs test (2 groups) or Friedman test (> 2 groups) followed by uncorrected Dunn’s multiple comparison test. Correlations were assessed using the Spearman’s rank (r_s_) correlation test. The tests used in each one of the graphs are referred and detailed on the figure legends. Analysis of the outliers was performed using the ROUT method (Q=0.1 %) and a maximum of three data points were removed. Statistical significance was achieved when *p*<0.05 (* *p*<0.05; ** *p*<0.01 and *** *p*<0.001).

## Results

### *DLEU1* and miR-16-5p expression levels are dysregulated during osteoclastogenic differentiation and correlate with osteoclastogenic markers

To identify new potential therapeutic targets for bone diseases caused by increased resorption activity, such as osteoporosis, the expression of the ncRNAs located in the chr13q14.2 region, specifically *DLEU1* / *DLEU2* / miR-15a / miR-16-1 (Supplemental Figure 1), was evaluated during the 9 days of osteoclastogenic differentiation from six human blood donors. Results show that *DLEU1* is significantly overexpressed at days 7 and 9, while *DLEU2* is downregulated since day 2 (Figure 1a). To validate these results, monocytes isolated from seven additional independent donors were cultured using a different protocol, with extended periods of osteoclastogenesis. Results confirm that *DLEU1* is significantly upregulated during differentiation, but could not confirm the observed downregulation of *DLEU2* (compare Figure 1a with Supplemental Figure 2). Importantly, only *DLEU1,* but not *DLEU2*, positively correlate with the osteoclastogenic markers *ACP5* (TRAcP5 encoding gene) (*r_s_*=0.569; *p*=0.037) and *CTSK* (*r_s_* =0.736; *p*=0.004) (Figure 1b), at day 9 of differentiation. Regarding the miRNAs, they were found to be downregulated: miR-15a-5p was significantly downregulated at days 2 and 7 of differentiation, whereas miR-16-5p was consistently decreased at days 7 and 9 (Figure 1a). Notably, the expression of miR-16-5p, but not miR-15a-5p, negatively correlates to *ACP5* expression (*r_s_*=-0.543; *p*=0.048), suggesting a negative regulation of osteoclastogenic differentiation (Figure 1b). Considering these results, *DLEU1* and miR-16 were selected for further assessment of their impact on the multinucleation process, cell fusion and bone resorption processes of OCs.

**Figure 1.**
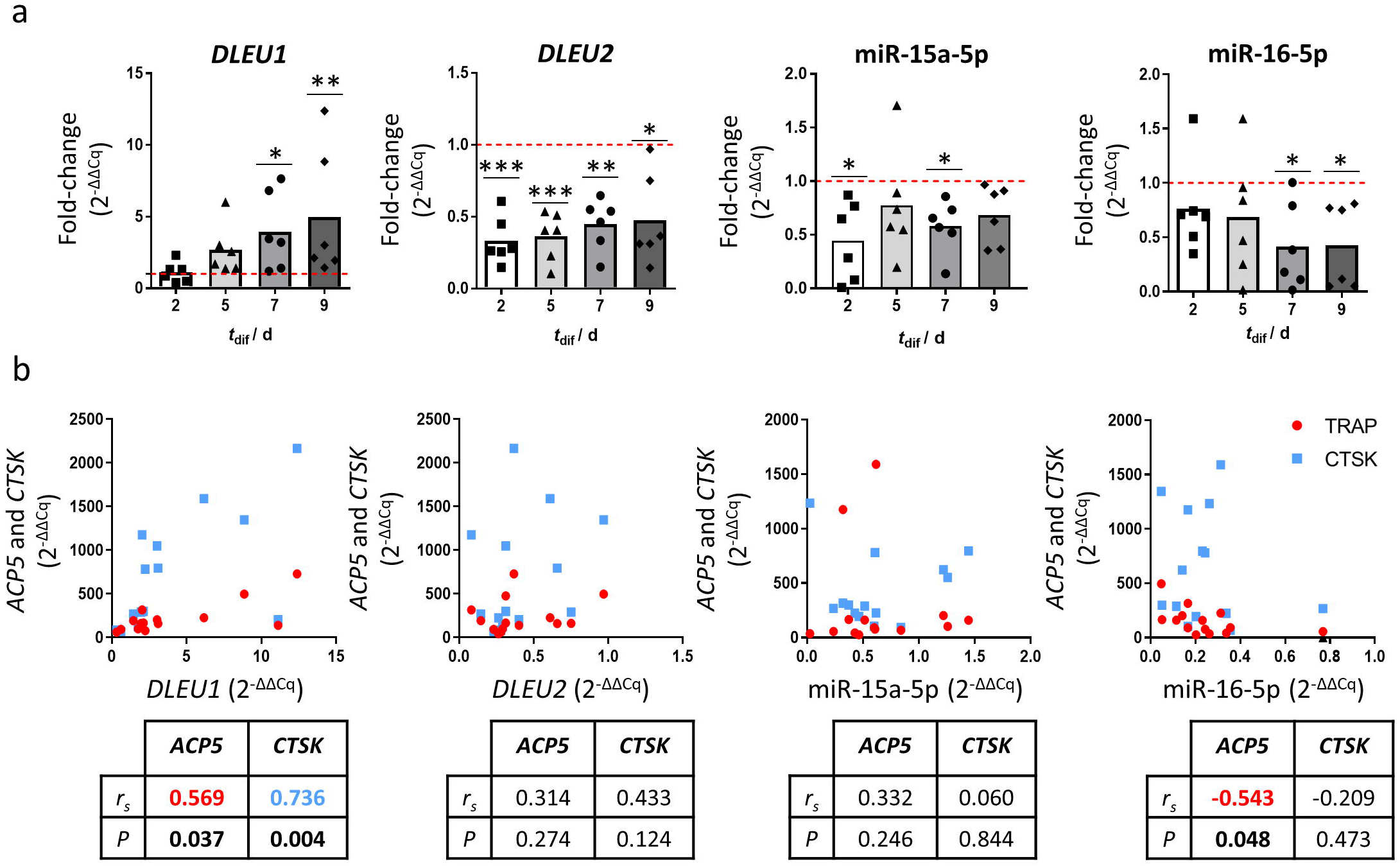
*DLEU1*, *DLEU2*, miR-15a-5p and miR-16-5p expression levels during osteoclastogenic differentiation of human primary OCs. Human primary monocytes isolated from old males (≥50 years old) buffy coats using the BD Imag Anti-Human CD14 Magnetic Particles kit were cultured for 2 days with 25 ng/mL of M-CSF (t_dif_: day 2) and 7 additional days with 25 ng/mL of M-CSF and 25 ng/mL of RANKL (t_dif_: day 5, 7 and 9). Expression levels are shown as a fold-change of day 0 levels (isolation day). **a)** *DLEU1*, *DLEU2*, miR-15a-5p and miR-16-5p expression levels during osteoclastogenic differentiation (*N*=6). **b)** Correlation of the expression levels of *DLEU1*, *DLEU2*, miR-15a-5p and miR-16-5p with osteoclastogenic markers (*ACP5* and *CTSK*) (*N*=14). Each dot represents the expression levels obtained from OCs generated *in vitro* from each individual donor. For each scatterplot, the Spearman correlation coefficient and significance levels are shown. (red: *ACP5* and blue: *CTSK*).

### Silencing of *DLEU1* and overexpression of miR-16 impair OCs’ fusion through distinct mechanisms

To investigate the potential involvement of *DLEU1* and miR-16 on OC fusion, human pre-OCs were transfected either with a silencing RNA against *DLEU1* (siDLEU1-OC) or with a miR-16 mimics (miR-16-OC), at day 3 of the differentiation process (Figure 2a). Results disclose a significant downregulation of *DLEU1* and an increase of miR-16, confirming a successful transfection (Supplemental Figure 3). Cell fractionation shows that *DLEU1* transcript, contrary to other typically nuclear lncRNAs, such as *XIST* and *MALAT1*, is mainly located in the cytoplasm (Supplemental Figure 4).

**Figure 2.**
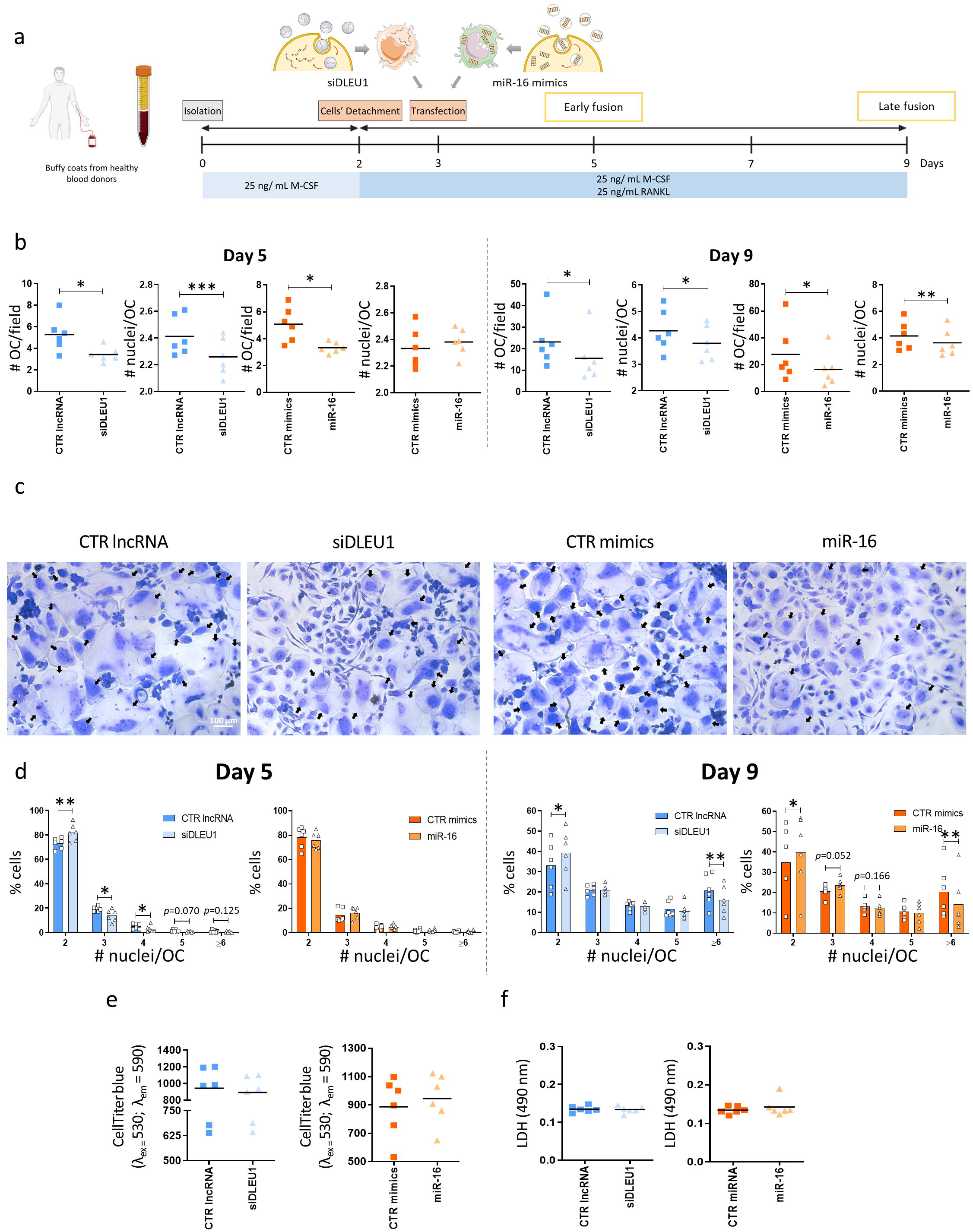
Impact of *DLEU1* and miR-16 on multinuclearity of human primary OCs. a) Experimental setup used to assess the impact on the fusion capacity and multinucleation; siDLEU1-OCs: OCs transfected with a siRNA against *DLEU1* and miR-16-OCs: OCs transfected with miR-16 mimics. **b)** Impact on the number of OCs per field (# OC / field) and number of nuclei per OC (# nuclei / OC) (*N*=6), at day 5 and 9 of the differentiation process. Each dot represents the mean of a minimum of 8 wells per condition (7 fields per each well) and donor. c) Representative images of OCs transfected with CTR lncRNA, siDLEU1, CTR mimics or miR-16 mimics at day 3 and left to differentiate until day 9. Black arrows are highlighting the multinucleated OCs. **d)** Histogram with the distribution of the percentage of the number of OCs with a specific number of nuclei at early and late fusion stages (*N*=6). Each dot represents the mean of a minimum of 8 wells per condition (7 fields per each well) and donor. **e)** Measurements of the metabolic activity through CellTiter-blue assay (*N* = 6). **f)** Quantification of LDH levels on the conditioned-media at day 5, 2 days after the transfection (*N* = 6).

Silencing of *DLEU1* and transfection with miR-16 mimics led to an impairment in the number of OCs formed per field (# OC/field), both at day 5 and 9 of differentiation (Figure 2b and 2c). Accordingly, this inhibition is consistently observed within the experimental replicates of each donor (Supplemental Figure 5). When the number of nuclei per OC (# nuclei/OC) was analyzed, the results evidenced a significant decrease for siDLEU1-OCs at day 5 and 9. Regarding miR-16-OCs, the # nuclei/OC was not impacted at early stages (day 5), with differences only being noticed at later stages (day 9), indicating a stage-specific regulation (Figure 2b). A detailed analysis of the abundance of OCs with a specific number of nuclei revealed a predominance of bi-nucleated OCs, regardless the condition, compared with OCs with more than 3 nuclei, at day 5, as expected for early stages of osteoclastogenesis (Figure 2d). Inhibition of *DLEU1* resulted in an enrichment in bi-nucleated OCs in detriment of more mature osteoclasts (with 3 and 4 nuclei) (day 5, Figure 2d), which is in line with the decrease observed in the # nuclei/OC at the same time-point (day 5, Figure 2b). In contrast, no statistical differences were observed after transfection of the pre-OCs with miR-16 mimics, when compared to the control condition (at day 5, Figure 2d). Consistent with the findings observed at day 9 (Figure 2b) regarding the # nuclei/OC, results show that both silencing of *DLEU1* and miR-16 mimics mediate cell fusion primarily through an increase of OCs containing 2 nuclei, in detriment of a marked decrease of more matured OCs (with six or more nuclei) (Figure 2d).

To test a potential synergistic effect on multi-nuclearity between downregulation of *DLEU1* and increased levels of miR-16, pre-OCs were co-transfected with both siDLEU1 and miR-16 mimics. Using pre-OCs from the same donor, a robust decrease of the # nuclei/OC and of the # OC/field was observed, for both end-points (days 5 and 9; Supplemental Figure 6a and 6b). This solid decline was consistently observed using cells from three different donors. As expected, the co-transfection with siDLEU1 and miR-16 mimics resulted in a mixture of the phenotypes, rather than a synergic effect obtained when pre-OCs were transfected separately, with either siDLEU1 or miR-16 mimics (compare Supplemental Figure 6a with Figure 2b). Moreover, results showed that the metabolic activity did not seem to be affected by siDLEU1 or miR-16 mimics (Figure 2e). Also, no differences in the LDH levels released into the conditioned-media were found between siDLEU1, miR-16 mimics and the controls, excluding a cytotoxic effect (Figure 2f).

### Proteome profiling unveil that *DLEU1* and miR-16 affect OCs fusion through independent mechanisms at early time-points

To screen for specific proteins regulated by *DLEU1* and miR-16, a mass spectrometry-based quantitative proteomic analysis was performed in siDLEU1- and miR-16-OCs at early stages of osteoclastogenesis (day 5, see Figure 2a). When considering a minimum requirement of two unique peptides per detected protein, a total of 931 proteins were identified (Figure 3a). Results show that, when compared with the respective controls, 25 proteins (16 upregulated and 9 downregulated) were differentially expressed in siDLEU1-OCs (*p*<0.05, Figure 3a and 3b), whereas 28 (25 upregulated and 3 downregulated) were significantly impacted in miR-16-OCs (*p*<0.05, Figure 3a and 3b). From the differentially expressed proteins, only 2 were simultaneously impacted by the silencing of *DLEU1* and miR-16 mimics (Supplemental Figure 7), namely Epoxide Hydrolase 1 (EPHX1; P07099) and Inter-alpha-trypsin Inhibitor Heavy chain H3 (ITIH3; Q06033). EPHX1 showed an opposite tendency between conditions, while ITIH3 is upregulated in both, suggesting that *DLEU1* and miR-16 act on OC differentiation through independent mechanisms.

**Figure 3.**
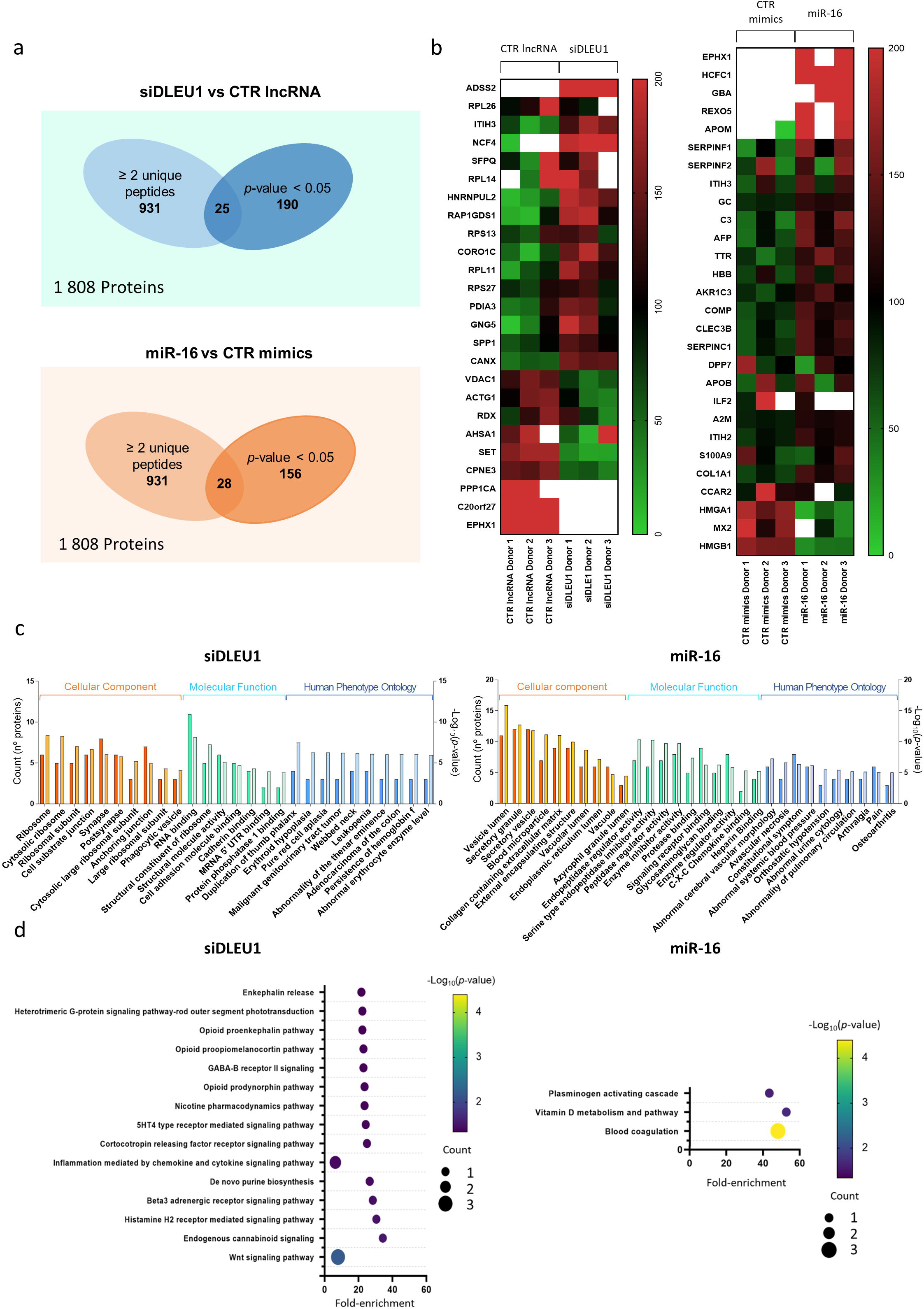
Proteomic profiling of *DLEU1* and miR-16 downstream targets and processes. **a)** Venn diagrams with the proteins differentially expressed after transfection with siDLEU1 and miR-16 mimics. **b)** Heatmaps with the differentially expressed proteins at day 5, with ≥ 2 unique peptides from three different donors after silencing *DLEU1* and overexpression of miR-16. **c)** GSEA software (Gene Set Enrichment Analysis) of the differentially expressed proteins. The x-axis shows the number of proteins and the y-axis the -Log_10_(*p*-value). **d)** Pathways predicted to be enriched when considering the differentially expressed proteins using PANTHER.

An enrichment analysis was performed using only the differentially expressed proteins to evaluate the processes and functions impacted. For siDLEU1-OCs, the differentially expressed proteins were associated with ribosomal complexes (such as “Ribosome”, “Cytosolic ribosome”, “Ribosomal subunit” and “Large ribosomal subunit”) and “Phagocytic vesicle” (Cellular Components); “Cell adhesion molecule binding” and “Cadherin binding” (Molecular Function); and numerous blood malignancies (Human Phenotype Ontology) (Figure 3c). On the other hand, when considering miR-16-OCs, the differently expressed proteins are related to the secretory activity, such as “Secretory granule”; “Secretory vesicle”; “External encapsulating structure” and “Vacuole” (Cellular Component); “Signaling receptor binding”, “Protease binding”, “Glycosaminoglycan binding” and “C-X-C Chemokine binding” (Molecular Function); and osteoarticular diseases, including Arthralgia and Osteoarthritis (Human Phenotype Ontology) (Figure 3c).

Interestingly, 7.9 % of these proteins are associated with the “Wnt signaling pathway” (P00057; siDLEU1; Figure 3d) and 12.5 % with the “Vitamin D metabolism and pathway” (P04396; miR-16; Figure 3d). Notably, proteomic results also show that depletion of *DLEU1* led to an overexpression of the Neutrophil Cytosolic Factor 4 (NFC4) protein, which is linked to the “*osteoclast differentiation”* pathway in Kyoto Encyclopedia of Genes and Genomes – KEGG (I04380; Supplemental Figure 8), and an impairment of the Actin Gamma 1 (ACTG1) protein, which is one of the proteins responsible for phagocytosis and formation of the phagocytic cup in the “*phagosome*” pathway (I04145; Supplemental Figure 8).

### Time-lapse recordings detail the impact of *DLEU1* and miR-16 on the OC fusion type and fusion pairs

Time-lapse analysis was used to track and follow the behavior of OCs at the single cell level, throughout 24 h, over the course of 4 days using cells from four different donors (Figure 4a). In line with the results obtained with the May-Grünwald staining (Figure 2c and 2d), a significant decrease in the fusion events was observed in siDLEU1- and miR-16-OCs (Figure 4b). Nonetheless, no differences were detected regarding the time at which the events occurred (Figure 4b). The fusion modalities of the OCs were also investigated (Figure 4c, 4d and Supplemental Figure 9). Regarding the fusion phenotype, the results showed a consistent decrease in the *phagocytic cup* mode (*Pha.Cup*) for siDLEU1-OCs, in comparison to the control, for the four donors used (Figure 4c and 4d), while an increase in the *from top* fusion mode was observed in three out of the four donors (Figure 4c and Supplemental Figure 9). However, no differences were found in the fusion phenotype when comparing miR-16-OCs with the control. Concerning the multi-nuclearity of the fusion partners, siDLEU1-OCs exhibit a specific impairment of fusion between mono and multinucleated OC, whilst there were no differences for miR-16-OCs (Figure 4e and Supplemental Figure 10).

**Figure 4.**
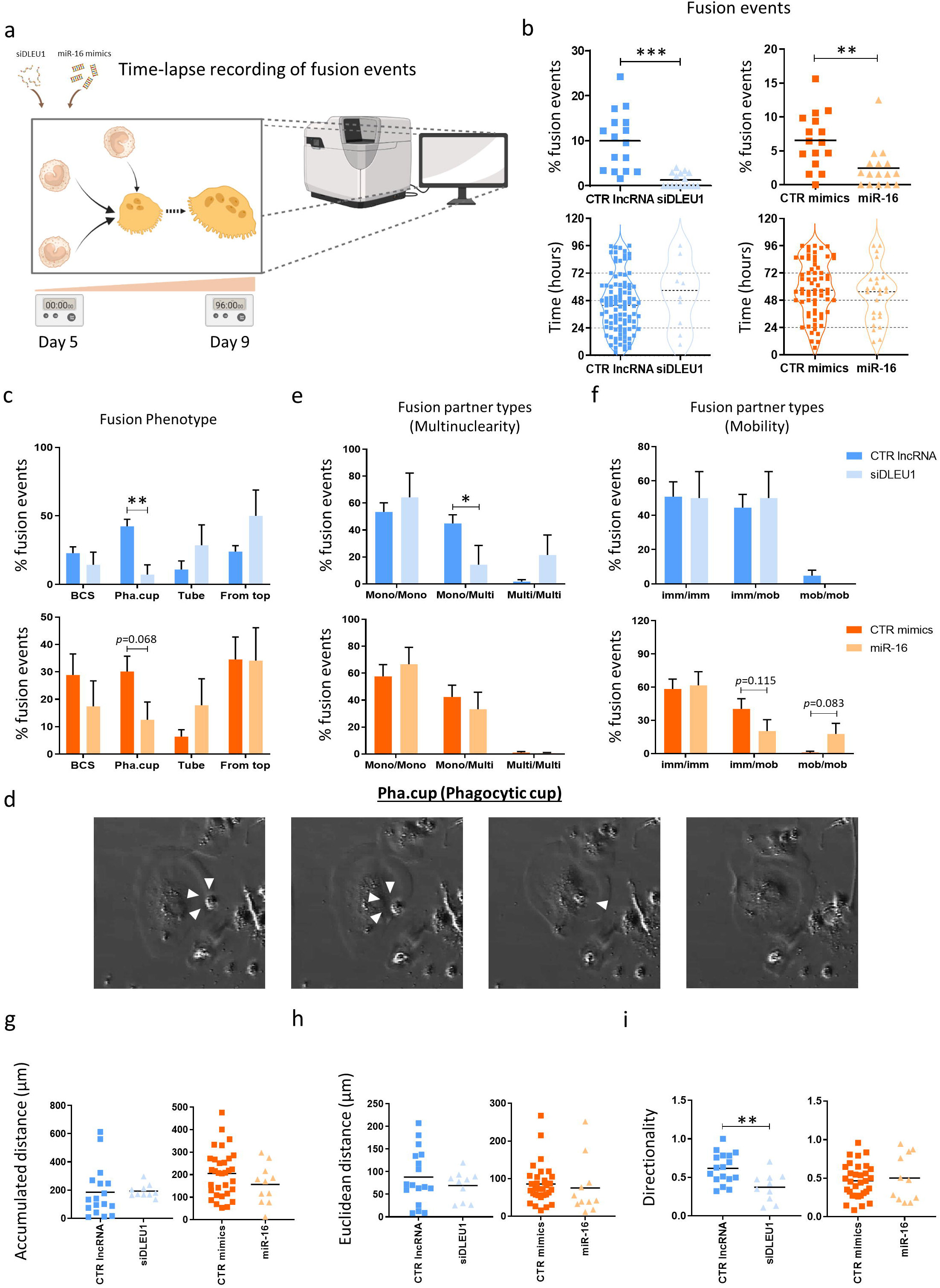
Time-lapse analysis of the cells undergoing fusion and respective characterization of the fusion events. **a)** Schematic illustration of the time-lapse recordings performed to assess the impact on cell fusion. **b)** Percentage of OCs performing fusion events and respective time they occur. **c)** Analysis of the frequency of the fusion events with a specific fusion modality and **d)** representative images of the phagocytic cup (*Pha.cup*) fusion modality. **e)** Analysis of the frequency of the fusion partners classified according to their nuclearity (mononuclear cells with mononuclear cells: mono-mono; mononuclear cells with multinucleated cells: mono-multi; multinucleated cells with multinucleated cells: multi-multi). **f)** Analysis of the frequency of the fusion partners classified according to their mobility. **g)** Accumulated distance; **h)** Euclidean distance and **i)** Directionality of the mobile OCs undergoing fusion. The graphs provided depict data from 16 different fields from a single donor, representative of a total of four donors.

Since the fusion capacity of the OCs can be compromised if cell mobility is affected, this parameter was also investigated for the OCs that fuse. No significant differences were found for either siDLEU1- or miR-16-OCs. However there is a tendency for a reduction of the fusion between immobile and mobile fusion partners (imm/mob), and an increase in the frequency of fusion between mobile and mobile pairs (mob/mob) for miR-16-OCs (Figure 4f). Furthermore, the accumulated travelled distance (Figure 4g), the euclidean distance (Figure 4h), and the directionality (Figure 4i) were measured in OCs at early stages of differentiation (day 5). Overall, no differences were observed for any of these parameters (Figure 4g and 4h), except for the directionality in siDLEU1-OCs, which travelled less linearly (Figure 4i). Overall, these results suggest that siDLEU1, but not miR-16, has an impact on OC fusion types, fusion partners and directionality of movement.

### Effect of *DLEU1* and miR-16 on the migratory capacity of the OCs

A comprehensive analysis encompassing all OCs (fusing and non-fusing) was then carried out, instead of solely focusing on OCs undergoing fusion, as performed in the previous section. Interestingly, results showed a lower percentage of immobile cells and an increase of OCs in mobile regime in the miR-16-OCs group, when compared with CTR mimics (5a and 5b). No differences were found for siDLEU1 (Figure 5a and 5b). Additionally, an increase in the accumulated distance (Figure 5c) and in the euclidean distance (Figure 5d) was observed for miR-16-OCs, while no differences were observed for siDLEU1-OCs (Figure 5c, 5d and 5e). A more general approach demonstrated that the shape parameters of the OCs were not altered by the modulation of *DLEU1* or miR-16 levels (Figure 5f). Moreover, in agreement with the LDH measurements, no differences were detected regarding the number of cells dying (Figure 5g). Altogether, the results suggest that miR-16 increases OC mobility, leading to enhanced migration (both accumulated and euclidean distance), compared with the control group (Figure 5a, 5c and 5d). These findings may indicate that since miR-16-OCs fuse less, the cells need to search more for the respective fusion partner (Figure 4b).

**Figure 5.**
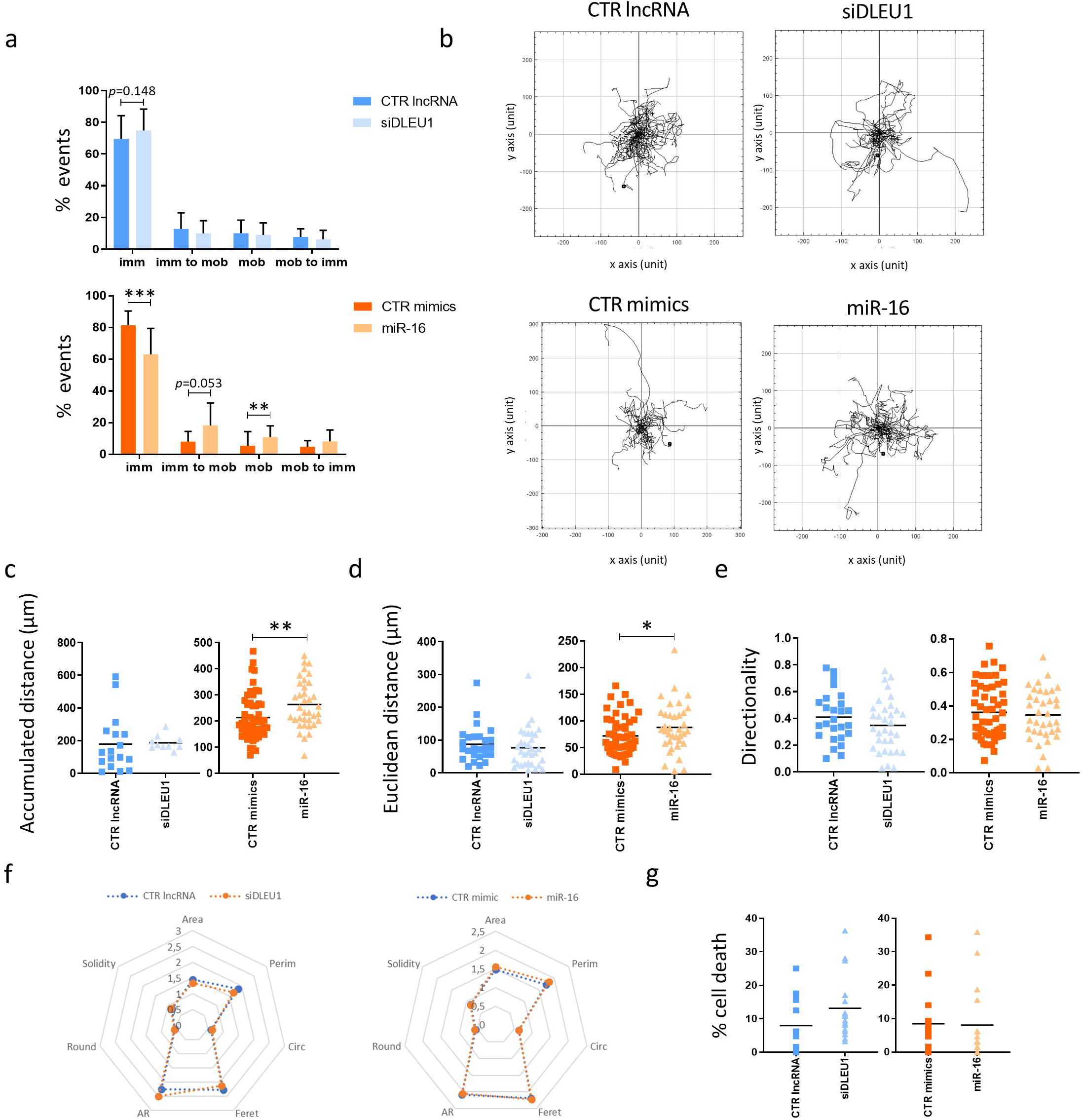
Time-lapse analysis of the migratory capacity, cell death and division of non-fusing and fusing cells. **a)** Analysis of the frequency of cells classified according to a specific migration pattern. **b)** Representative track of mobile OCs per condition. **c)** Accumulated distance; **d)** Euclidean distance and **e)** Directionality of the mobile OCs. **f)** Cell area / shape parameters and; **g)** Percentage of cells dying. The graphs provided depict data from 16 different fields from a single donor, representative of a total of four donors.

### Differential impact of *DLEU1* and miR-16 on Resorption Modes

We next investigated whether *DLEU1* and/or miR-16 affect bone resorption. The expression profile of *DLEU1* and miR-16 was evaluated 72 h after seeding mature OCs onto bone slices and allowing them to resorb bone. Results revealed a significant reduction in the miR-16 expression levels for all six donors, while the expression of *DLEU1* exhibited heterogeneity and non-statistically significant differences, with five out of the six donors showing an upregulation (Supplemental Figure 11). These findings suggest a stronger impact of miR-16 mimics, over siDLEU1, on the bone resorption capacity. To further investigate this, mature OCs were transfected with siDLEU1, miR-16 mimics, or the respective controls, on day 8 of the differentiation process. The transfected OCs were then seeded on top of bone slices on the next day (day 9) and allowed to resorb bone until day 12 (Figure 6a).

**Figure 6.**
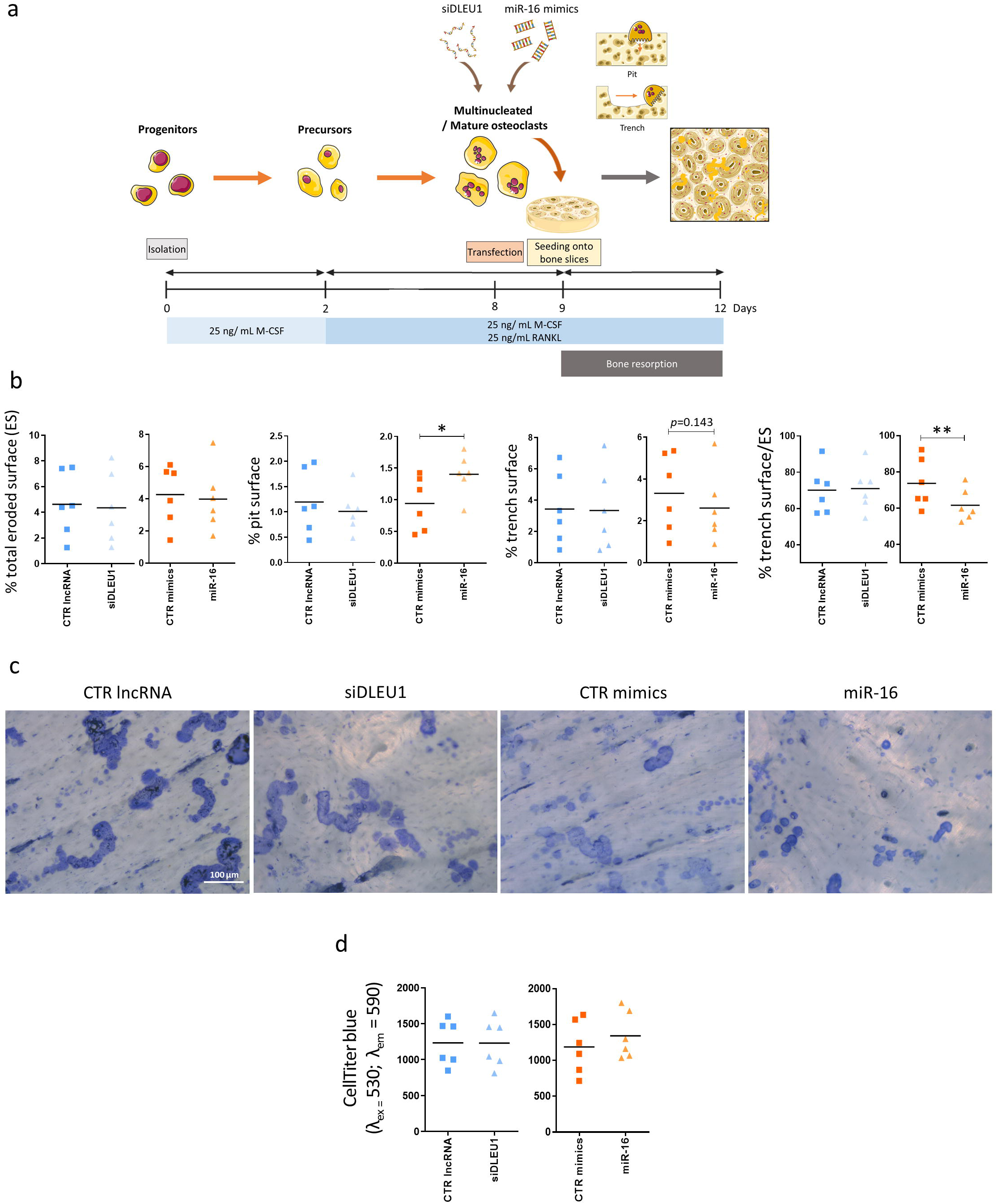
Impact of *DLEU1* and miR-16 on bone resorption. **a)** Scheme of the experimental design used to assess the impact on bone resorption. **b)** Measurement of the eroded surface by OCs after *DLEU1* silencing and miR-16 overexpression (*N*=6) (5 slices per condition and per donor). **c)** Representative images of the resorbed area from OCs transfected with CTR lncRNA, siDLEU1, CTR mimics or miR-16 mimics at day 8, seeded onto bone slices at day 9 and left to resorb until day 12 (5 slices per condition and per donor). **d)** Measurements of the metabolic activity through CellTiter-blue assay (*N* = 6) after 3 days on top of the bone slices.

End-point analyses of the bone slices at day 12 showed that neither the silencing of *DLEU1* nor the overexpression of miR-16 affected the percentage of total eroded surface (Figure 6b and 6c). For siDLEU1-OCs, no differences were detected regarding the resorption modalities (pits and trenches) (Figure 6b). In contrast, miR-16-OCs exhibited a shift in the resorption mode towards an increase of OCs forming pits, in detriment of those forming trenches, which is significantly decreased (Figure 6b). Finally, the metabolic activity was assessed at day 12, showing not to be affected neither for siDLEU1- nor miR-16-OCs (Figure 6d).

### Time-lapse recordings detail the impact of *DLEU1* and miR-16 on each resorption mode

To further elucidate the impact on parameters associated with the resorption mode of the OCs and understand the variations observed in the end-point analysis, we studied the behavior of siDLEU1- and miR-16-OCs over a 72 h period using cells from three independent donors. No significant differences were observed after silencing of *DLEU1* (Figure 6c and 7a). However, an increase in the number of OCs making pits and a reduction in those making trenches, were observed for miR-16-OCs, when compared with the respective control (Figure 7a). Regarding the OCs performing pits, no differences were observed regarding the resorption duration (Figure 7b) and for the diameter of the formed pits (Figure 7c), for all the conditions. The average pit area also remained unaffected (Figure 7d) by siDLEU1 or miR-16 mimics. On the other hand, siDLEU1 and miR-16 negatively impacted OCs’ resorption speed (Figure 7e) and the ability of a single OC to perform multiple pits (Figure 7e, 7f and 7g).

**Figure 7.**
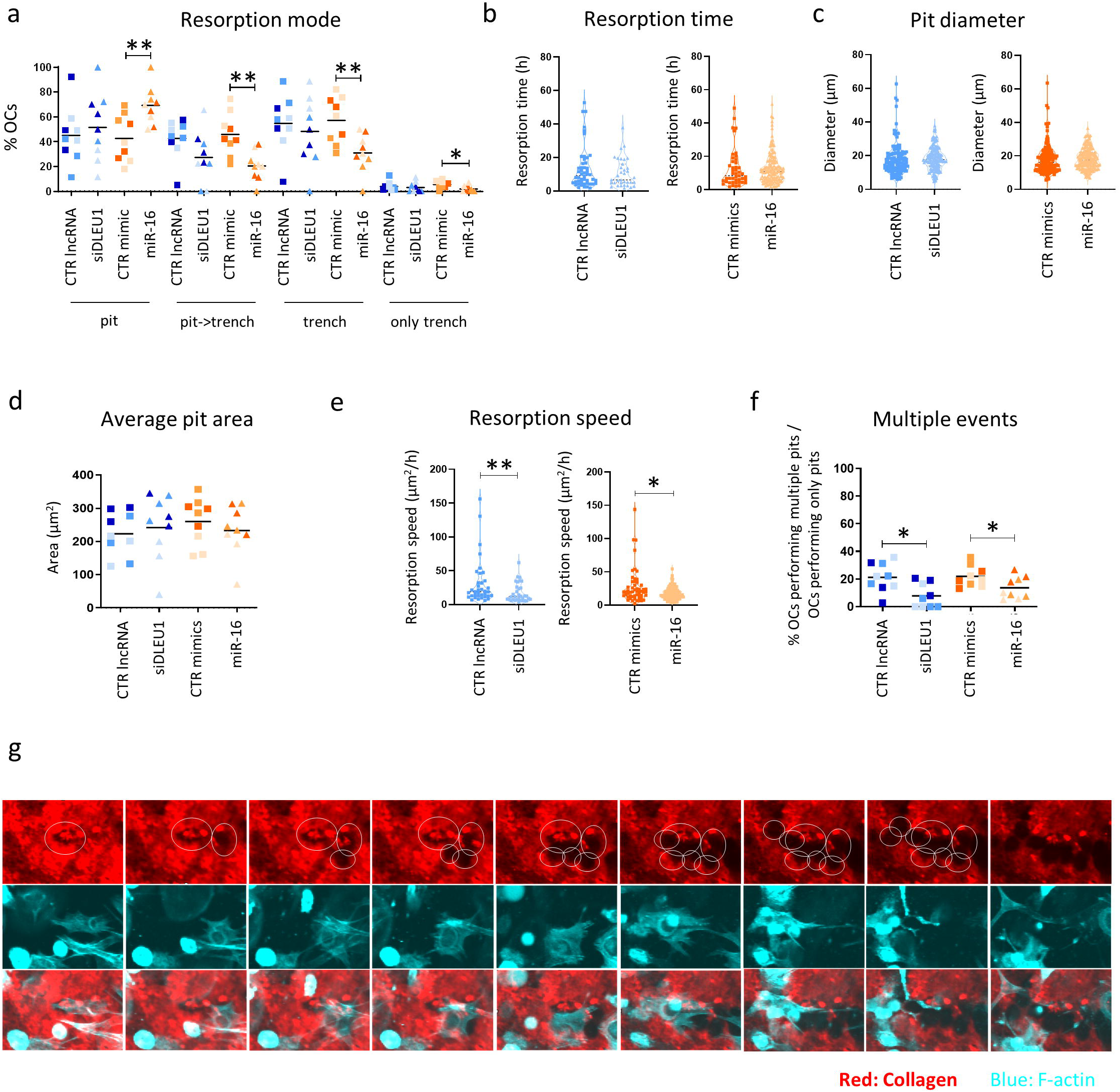
Time-lapse analysis of the resorption mode and resorptive capacity of OCs performing pits. **a)** Percentage of OCs categorized according to the resorption mode. Each dot represents the mean obtained in each video analyzed (3 per donor). For each condition dots with the same shades correspond to videos from the same donor. (OCs that only form pits: pit; OCs that form trenches that started as pit: pit->trench; OCs that form trenches without starting with a pit: only trench; OCs that form trenches, either starting with a pit or not: trench). **b)** Resorption time taken by the OCs to make a pit. The resorption time of each OC from the 3 recordings analyzed from a representative donor, doing a pit was assessed and is shown as a dot. **c)** Diameter of the pit cavities. The diameter of each pit from 3 recordings analyzed from a representative donor. **d)** Average pit area. Each dot represents the mean obtained in each video analyzed (3 per donor). For each condition dots with the same shades correspond to videos from the same donor. **e)** Resorption speed of the OCs when doing pits. The resorption speed of the OCs doing pits from 3 recordings analyzed from a representative donor. **f)** Percentage of OCs that perform multiple pits when considering only the pit-forming OCs. Each dot represents the mean obtained in each video analyzed (3 per donor). For each condition dots with the same shades correspond to videos from the same donor. **g)** Selected time-lapse images of OCs making multiple pits over the course of 72 h.

Finally, a series of analyses focusing on the impact of both ncRNAs on trench mode was conducted. Time-lapse results show no differences for siDLEU1-OCs, while in the miR-16-OCs group, a significant reduction in the area resorbed was observed for OCs making trenches (Figure 8a), corroborating the results obtained in the end-point analysis (Figure 6c). This reduction may be partly explained by the decrease in the average area resorbed per trench and by the negative impact on the resorption speed in the miR-16-OCs group, when compared with the control (Figure 8b, 8c and 8d). Despite being slower, miR-16-OCs still maintained their ability to perform multiple trenches during the time-lapse recordings (Figure 8e). Regardless of the parameters studied, no significant changes were detected after manipulating the levels of *DLEU1* (Figure 8a, 8b, 8c, 8d and 8f). Overall, these results indicate that the impact of siDLEU1 on bone resorption is mainly observed on OCs making pits, while miR-16 affects both pit and trench formation.

**Figure 8.**
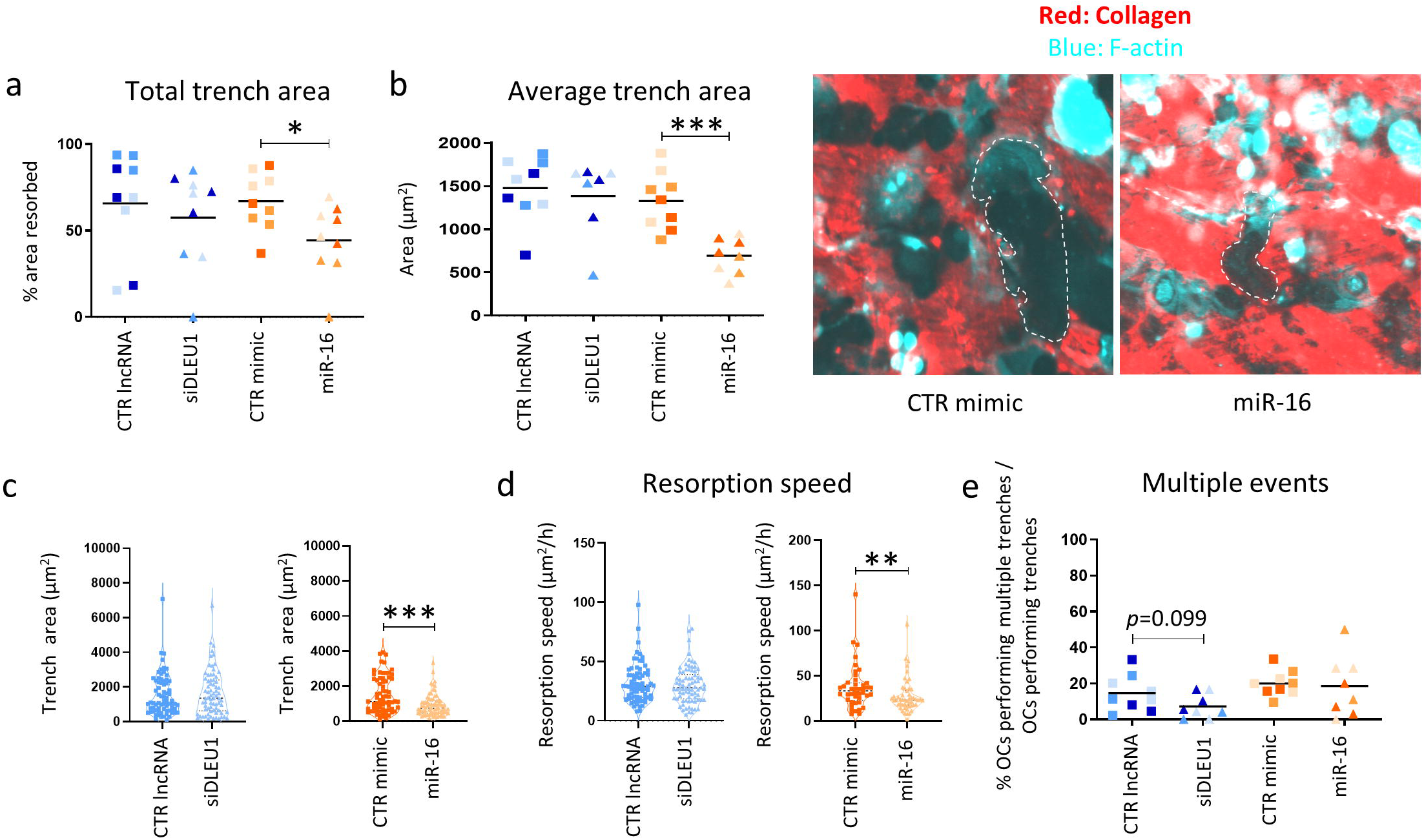
Time-lapse analysis of the resorptive capacity of OCs performing trenches. **a)** Percentage of area covered by trenches. Each dot represents the mean obtained in each video analyzed (3 per donor). For each condition dots with the same shades correspond to videos from the same donor. **b)** Average area of the trenches and representative time-lapse images. For each condition dots with the same shades correspond to videos from the same donor. **c)** The area of each trench from 3 recordings analyzed from a representative donor. **d)** Resorption speed of each trench from 3 recordings analyzed from a representative donor. **e)** Percentage of OCs that perform multiple trenches when considering only the trench-forming OCs. Each dot represents the mean obtained in each video analyzed (3 per donor). For each condition, dots with the same shades correspond to videos from the same donor.

## Discussion

Over the past two decades, the chromosomal region 13q14, housing the *DLEU1* and *DLEU2*/miR-15a/16-1 cluster, originally recognized as tumor suppressors [66,67], has emerged as a focal point of considerable attention in the context of several diseases. Notably, the discovery of deletions involving miR-15 and miR-16 in chronic lymphocytic leukemia (CLL) [68] marked a pivotal milestone that ignited a subsequent urge in research within the non-coding RNA (ncRNA) field. In bone biology, modulation of ncRNAs has been shown to promote bone regeneration [2], but their role in OC physiology and bone resorption remains far less explored. Presently, there are more than a dozen ongoing clinical trials exploring the therapeutic potential of miRNAs and siRNAs [69–72]. In contrast, the number of clinical trials involving lncRNAs remains limited, primarily due to a lack of comprehensive understanding of their functions [73]. In this study, we conducted a comparative analysis of the molecular functions of the lncRNA *DLEU1* and the small ncRNA miR-16-5p on OC biology. For this, we exclusively used human primary monocyte-derived OCs, deliberately avoiding the use of cell lines. This strategy effectively accounts for the natural variations between the different blood donors [16,18,74], thus providing a closer *ex-vivo* representation of the human *in vivo* settings. In this study, monocytes isolated from male donors were used due to the fact that OCs derived from males tend to exhibit a higher degree of aggressiveness, when compared to those derived from females [18,20,75], since we hypothesized that our treatments would hinder OC fusion and/or OC bone resorption capacity. Additionally, monocytes were obtained from older individuals since there is a notable increase in OC activity and higher levels of active CTSK in this demographic group [7,76].

To address the functional processes affected by the modulation of *DLEU1* and miR-16, we did not only perform end-point analyses, but also addressed the impact of modulating *DLEU1* and miR-16 on OCs at the single cell level by conducting time-lapse analyses. This enabled us to closely examine individual fusion events. Considering that multinucleation is not a random process and can be achieved through different fusion routes [30,32,64,77], this approach was fundamental in resolving the question of how a certain end-point result was attained. It also allowed to determine the involvement of fusion-related factors, which would otherwise be challenging to quantify. In our analysis we also included OCs with two nuclei to avoid missing the first fusion events. The absence of differences on the LDH levels and metabolic activity ruled out an effect on OC fusion due to an effect on cell viability.

The recordings revealed an impact on fusion modalities, in particular a reduction in the *Pha.cup* fusion mode in siDLEU1-OCs. This impairment may be partially attributed to the impact of siDLEU1 on the levels of the cytoskeletal protein ACTG1, linked to the term Phagosome (I04145), which includes the endocytic/phagocytic pathways, that are reported to be implicated in the fusion of OCs and macrophages [78,79] through mechanisms matching the *Pha.cup* mode. Interestingly, the impairment of endocytosis observed after *Actg1* knockout [80] also strengthens this hypothesis. Additionally, a decrease is observed in the fusion events between mono- and multinucleated partners when *DLEU1* levels are decreased. This reduction is consistent with the decreased number of OC with ≥3 nuclei, observed at both early and late stages of osteoclastogenesis. Together with the higher number of bi-nucleated OCs in the siDLEU1 condition, these results suggest an impairment of the fusion capacity through an obstruction in the fusion between smaller OCs. When compared to miR-16-OC, no differences in the multinucleation were found at early fusion stages, nevertheless an accumulation of bi-nucleated OC and a decrease in OC with more than 6 nuclei was observed at late fusion stages. Additionally, no differences were observed regarding the fusion partners and modes for the miR-16-OC group. These results collectively suggest that miR-16 specifically targets pre-OC, reducing their ability to fuse. Interestingly, despite the reduction in fusion events for miR-16-OC, the data revealed an increase in cell migration. While this may initially appear contradictory, as increased migration typically implies a greater potential for fusion, the higher migratory profile may indicate that cells need to search more extensively for their respective fusion partners. In a study on giant cell tumors, Sang *et* al. found that miR-16 promoted OC formation in mice bone marrow-derived macrophages [81]. Another miRNA from the same family, miR-195 [82], has also been reported to negatively regulate the expression of key osteoclastogenic genes [83]. These reports further support our findings, demonstrating that miR-16 impairs OC formation.

To support the potential of ncRNA as a new class of anti-resorption molecules, we conducted an in-depth investigation into the impact of miR-16 mimics and siDLEU1 on bone resorption, a unique feature of the OCs. Although these molecules do not impact the total area eroded by the OCs, they have a differential impact on the resorption mode, which has been shown to be clinically relevant and associated with the severity of OC resorption [7,16]. Specifically, Vanderoost *et* al. reported that for matching levels of erosion, the bone is more fragile when trenches are predominant [84]. Accordingly, OCs making trenches have higher erosion rates, create deeper cavities and have higher levels of active CTSK, reflecting a more aggressive form of resorption than pits [20,65]. Additionally, the importance of distinguishing resorption cavities between pit and trenches is even more evident when considering that the proportion of surface covered by trenches is increased by glucocorticoids [49] and inhibited by odanacatib [20], well known to elevate the bone mineral density in postmenopausal women with reduced bone mass [85]. In our findings, only miR-16-OCs, but not siDLEU1-OCs, exhibited a negative impact on the trench resorption mode, specifically concerning the average trench area and resorption speed. Regarding pit formation, while both siDLEU1 and miR-16 mimics influenced this resorption modality, the differences were more prominent for miR-16-resorbing-OC, culminating in an increase in the total eroded area covered by pits. Thus, delivery of miR-16 mimics into mature OCs seems a more promising approach to decrease the OCs aggressiveness, though not abolishing it.

The mass-spectrometry analysis provided further support for the hypothesis that *DLEU1* and miR-16, while sharing some functional overlap, operate through distinct mechanisms. Interestingly, only two proteins were found to be commonly impacted by both ncRNAs. Notably, several of the differentially expressed proteins have a role in regulating osteoclast physiology and activity. For example, An *et* al. identified that mitochondrial proteins, among them the voltage-dependent anion-selective channel protein 1 (VDAC1), are decisive for the formation of mature OCs and to have a pivotal function associated with the balance between OCs’ bone-resorbing activity and survival [86]. Treatment with an anti-VDAC antibody resulted in the inhibition of osteoclastogenesis and bone resorption of human OCs *in vitro* [87]. Accordingly, we observe an impairment of VDAC1 in siDLEU1-OCs. On the other hand, lysosomal acid glucosylceramidase (GBA), whose inhibition was previously shown to increase the number of nuclei per OC and OCs formation, as well as enhanced bone resorptive capacity [88], was identified by us to be one of the top upregulated proteins in miR-16-OC. Accordingly, Sibert *et* al. described miR-16-5p to positively regulate the GBA activity and expression in human fibroblasts [89]. Furthermore, several members of the serine protease inhibitor family, specifically SERPINF1, SERPINF2 and SERPINC1, exhibited high expression levels in miR-16-OCs. However, only SERPINF1, also known as PEDF, has been reported to negatively affect osteoclastogenesis, RANKL-mediated survival and bone resorption activity [90].

Overall, both ncRNAs *DLEU1* and miR-16 hold significant promise for the development of targeted treatments to counteract bone resorption and osteoporosis-related complications, either by influencing the fusion process and/or by directly impairing the bone resorptive activity of the OCs.

## Authors contributions

Conceptualization, S.R.M., K.S. and M.I.A.; methodology, S.R.M., A.B.S., J.B.O., M.A.B, K.S.; data collection, S.R.M., A.B.S.; statistical analysis, S.R.M., K.S. and M.I.A.; writing—original draft preparation, S.R.M., A.B.S., K.S., M.I.A.; writing-review and editing, S.R.M., A.B.S., J.B.O., M.A.B., K.S. and M.I.A.; funding acquisition, M.A.B. and M.I.A. All authors have read and agreed to the published version of the manuscript.

## Acknowledgements

The authors would like to acknowledge that this publication is under the scope of a collaboration formed between members (S.R.M., K.S. and M.I.A.) of the COST Action CA18139 GEMSTONE, supported by COST (European Cooperation in Science and Technology). COST is a funding agency for research and innovation networks.

The authors would like to thank to Cell Culture and Genotyping (CCGEN) and Proteomics (ROTEIRO/0028/2013; LISBOA-01-0145-FEDER-022125) services/platforms at i3S for technical support.

---

**Supplemental Figure 1.**
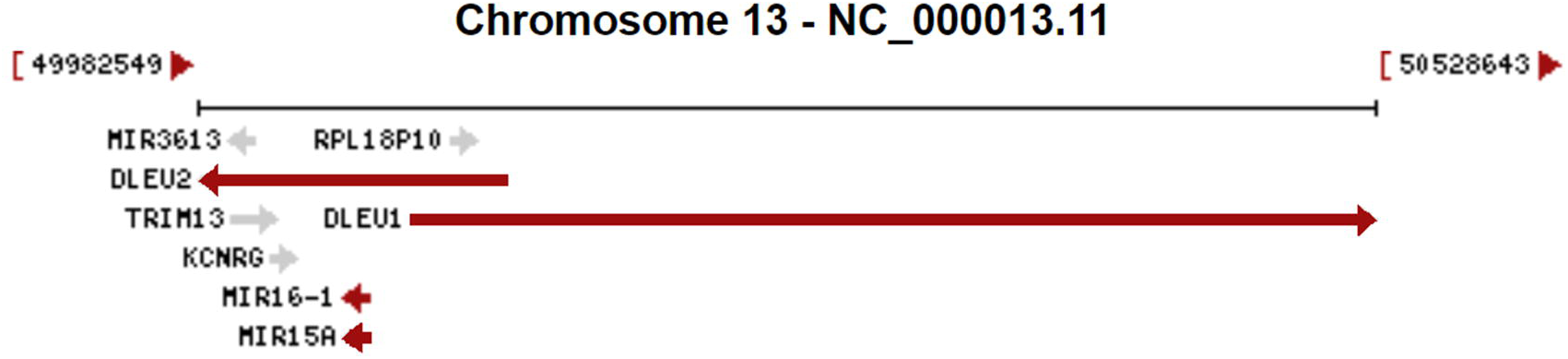
Schematic representation of the *DLEU1* / *DLEU2* / *miR-16-1*/*miR-15a* locus.

**Supplemental Figure 2.**
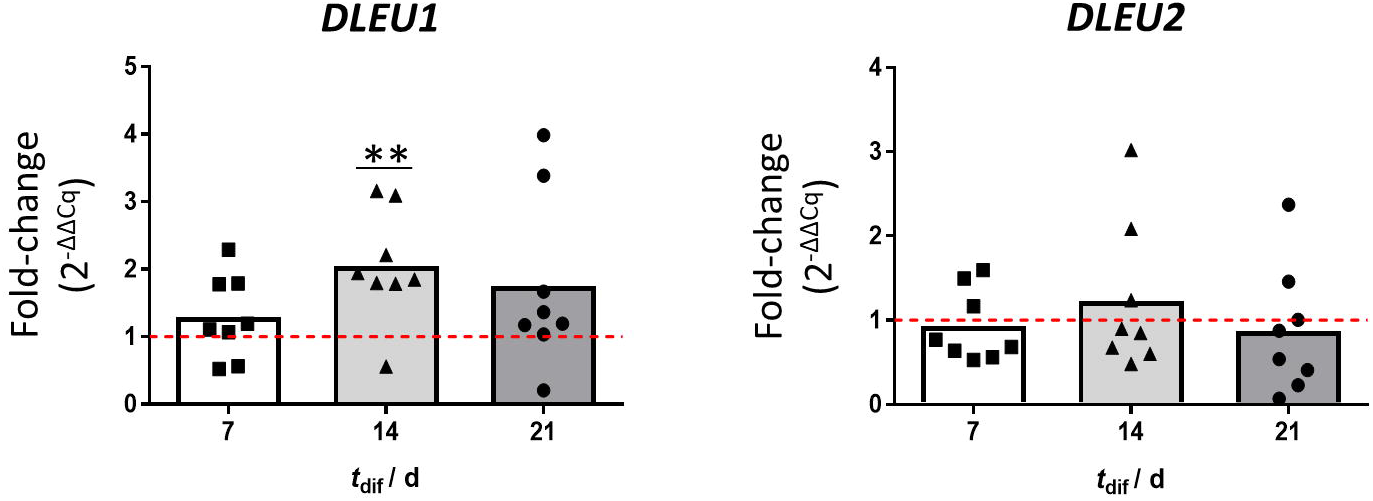
Expression levels of *DLEU1* and *DLEU2* during osteoclastogenic differentiation of human primary monocytes isolated using RosetteSep™ Human Monocyte Enrichment Cocktail (*N*=8), in comparison to day 0 (isolation day).

**Supplemental Figure 3.**
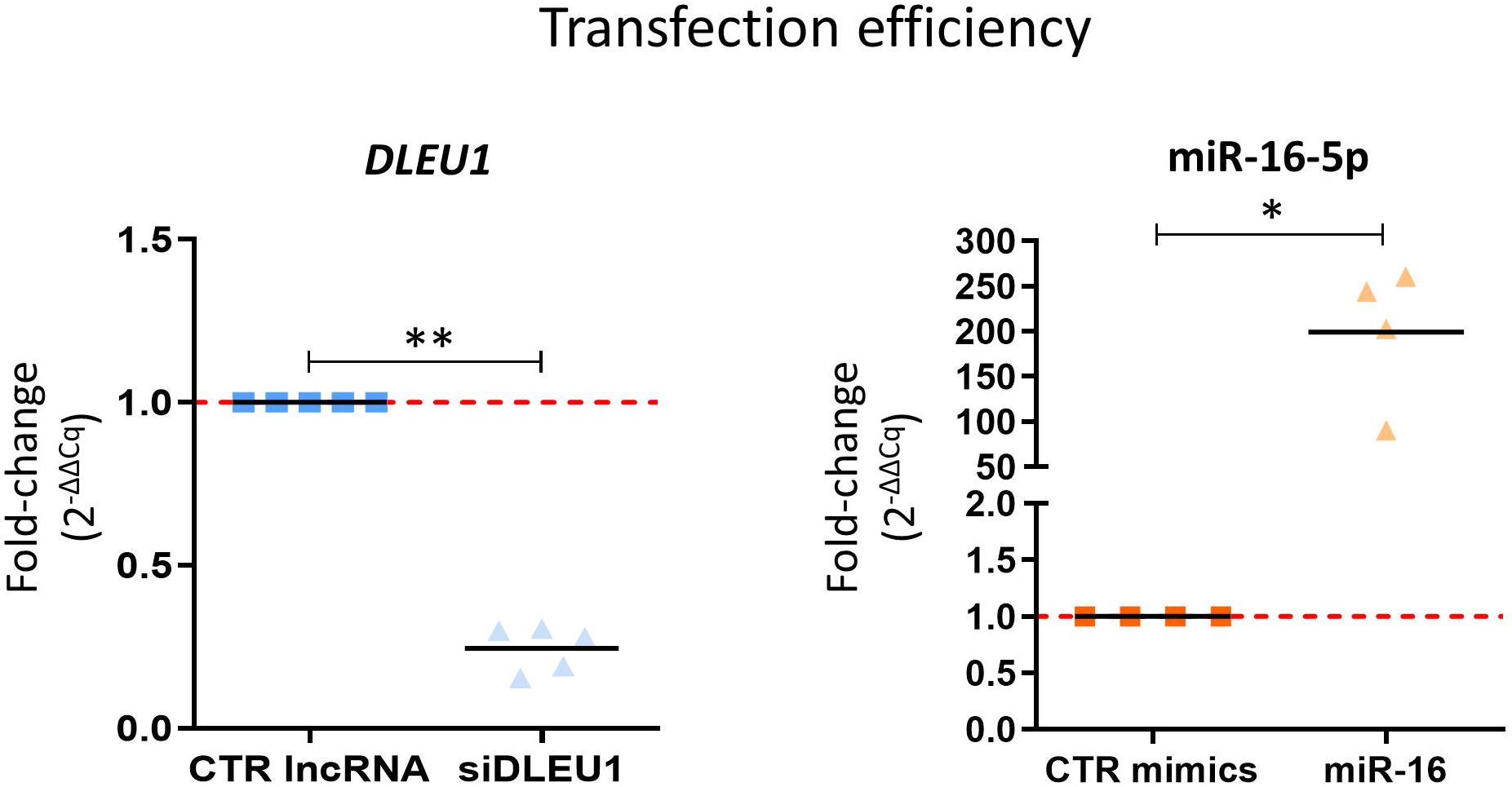
D*L*EU1 and miR-16 transcript levels (2^-ΔΔCq^) at day 5, in OCs after transfection at day 3 with siDLEU1, miR-16 mimics, or respective controls. Each dot represents a different donor.

**Supplemental Figure 4.**
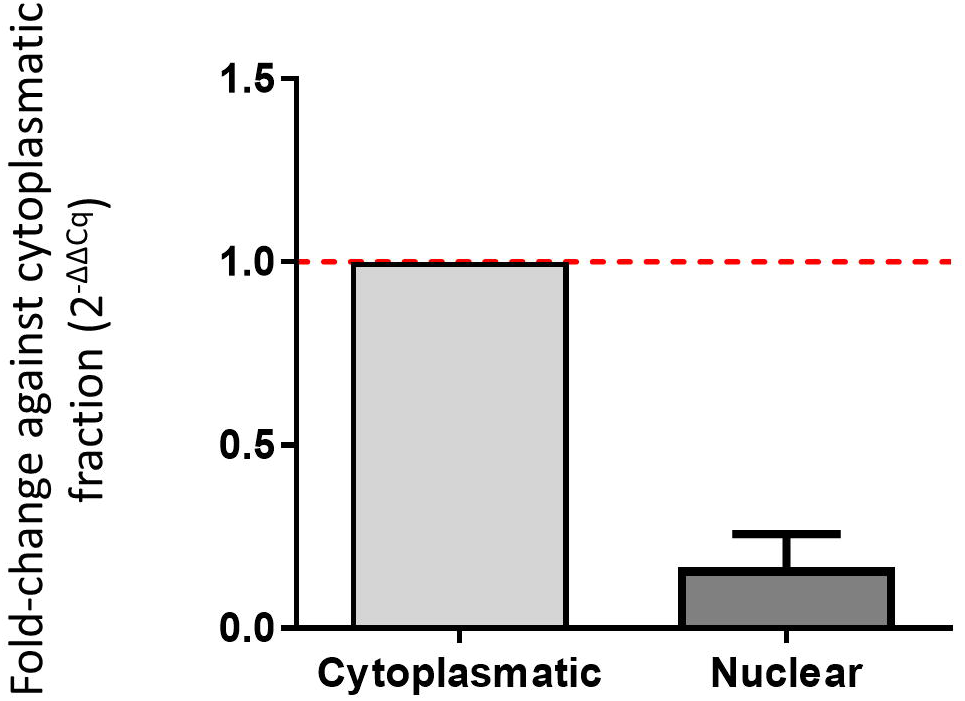
D*L*EU1 subcellular location in monocytes. Levels were normalized against the cytoplasmatic-cell content.

**Supplemental Figure 5.**
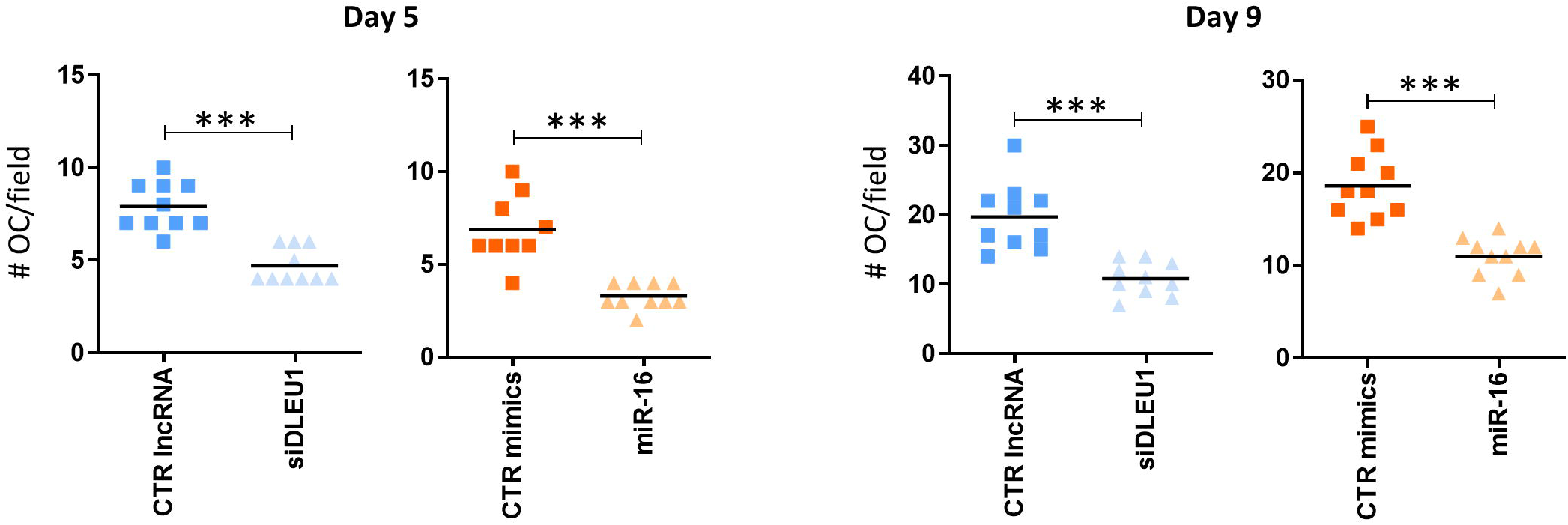
Representative distribution of the number of OC per field of one donor at day 5 and 9 of the differentiation process. Each dot represents the mean of OCs per each well (7 random fields counted per well).

**Supplemental Figure 6.**
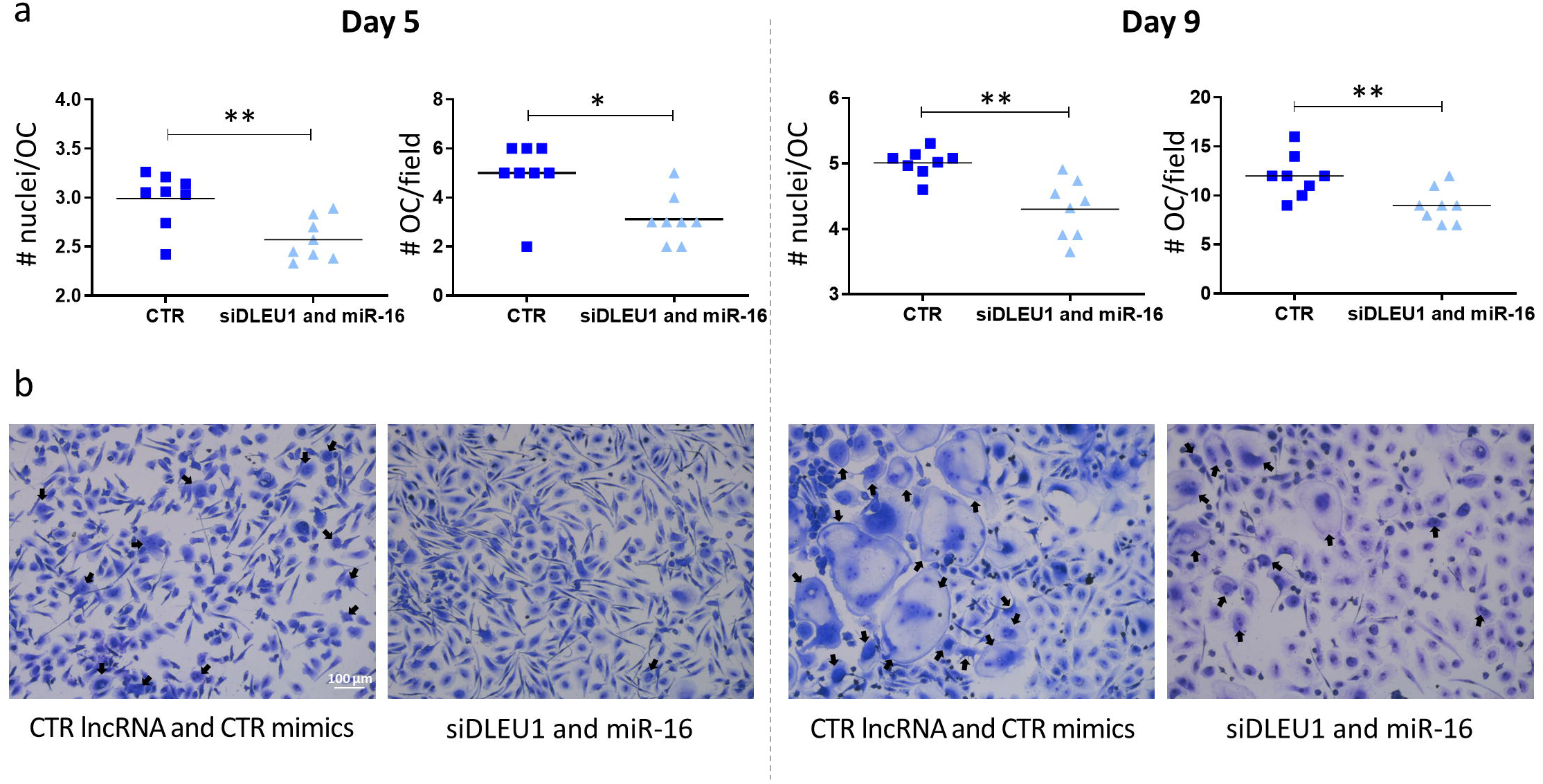
Impact of the co-transfection of *DLEU1* and miR-16 on multinuclearity. **a)** Impact on the number of nuclei per OC (# nuclei / OC) and number of OCs per field (# OC / field) (*N*=3), at day 5 and 9. Each dot represents the mean of OCs from 7 random fields per each well. **B)** Representative images of OCs co-transfected with siDLEU1 and miR-16 at day 3 and left to differentiate until day 5 and 9. Black arrows are highlighting the multinucleated OCs.

**Supplemental Figure 7.**
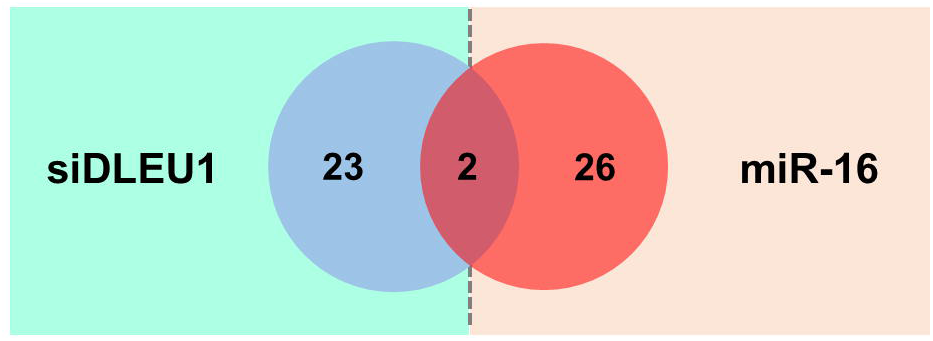
Venn diagram with the proteins differentially impacted by siDLEU1 and miR-16 mimics, simultaneously.

**Supplemental Figure 8.**
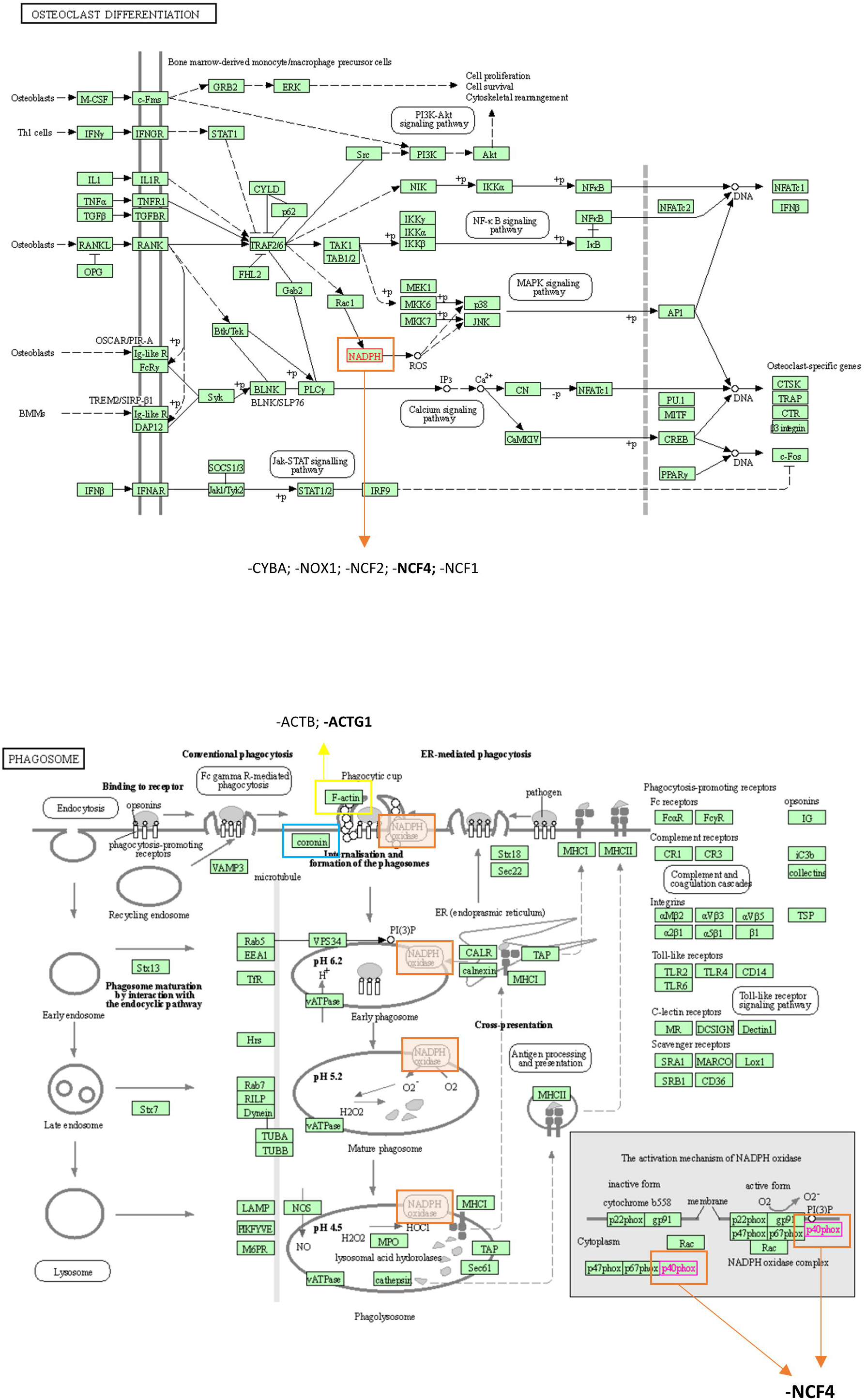
“Osteoclast differentiation” and “phagosome” are KEGG pathways predicted to be impacted in response to *DLEU1* silencing. Proteins discovered to be differentially expressed following proteomic analysis are highlighted in bold.

**Supplemental Figure 9.**
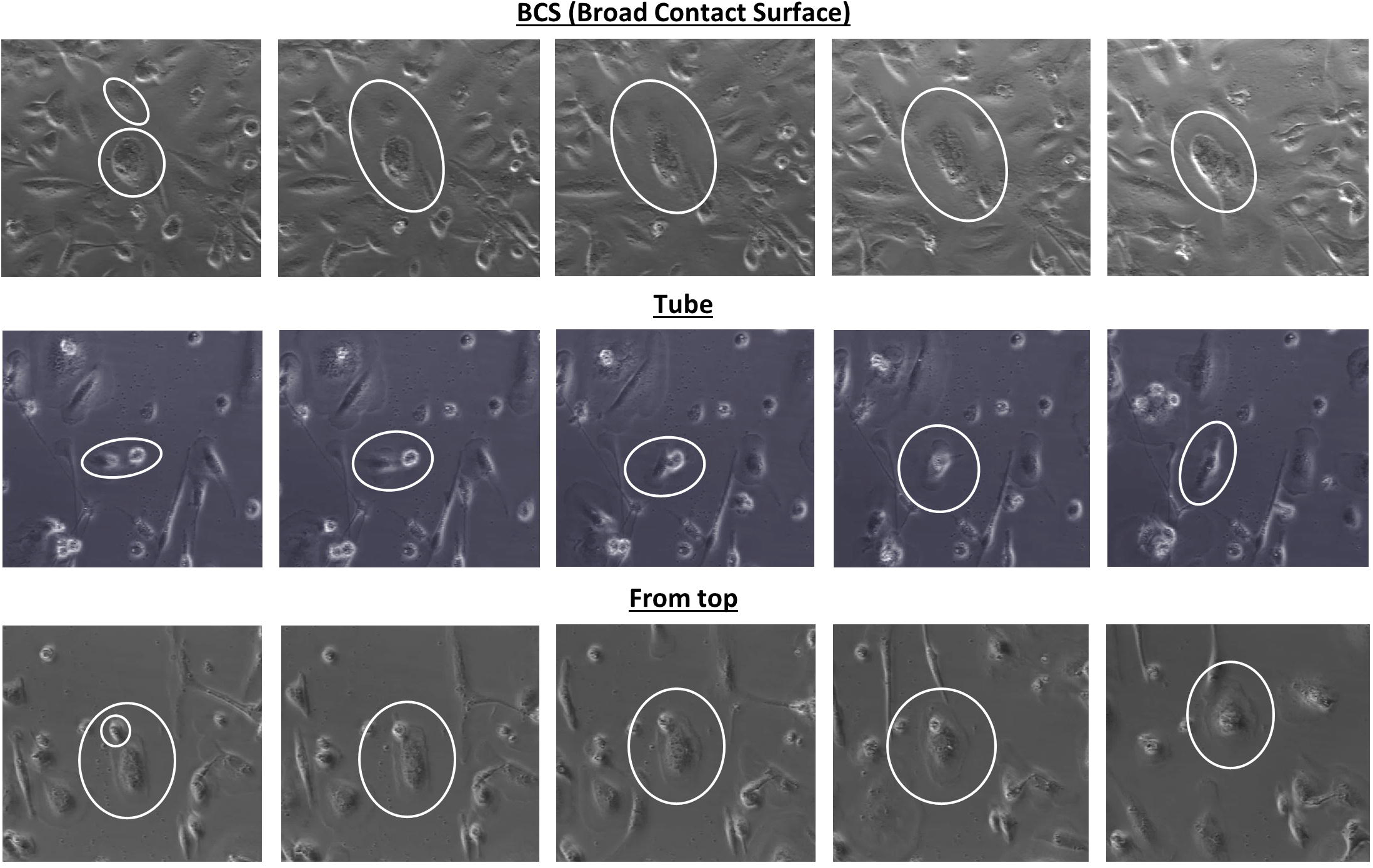
Representative images of the different types of fusion modalities used in our analyses.

**Supplemental Figure 10.**
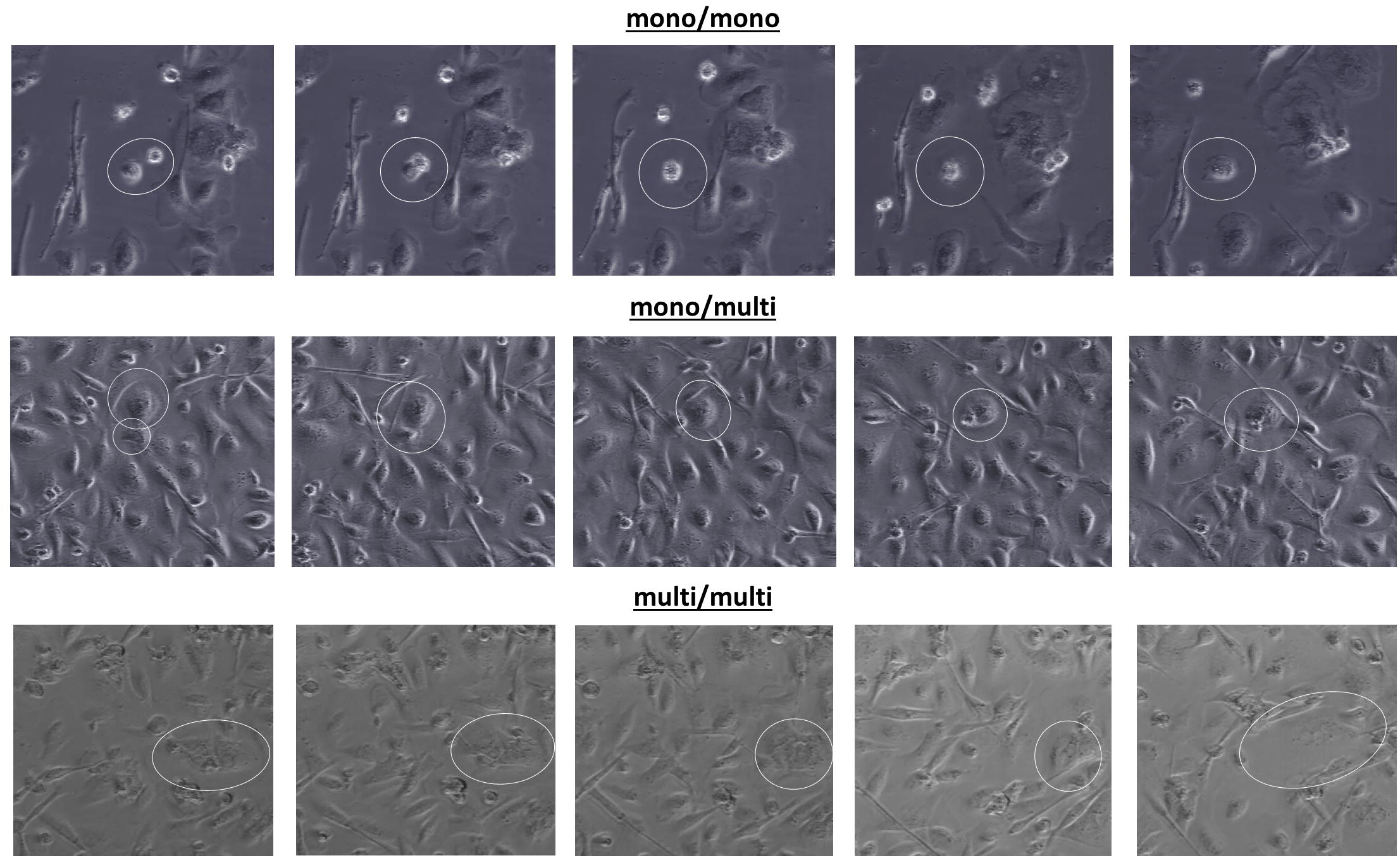
Representative images of the different types of fusion pairs used in our analyses.

**Supplemental Figure 11.**
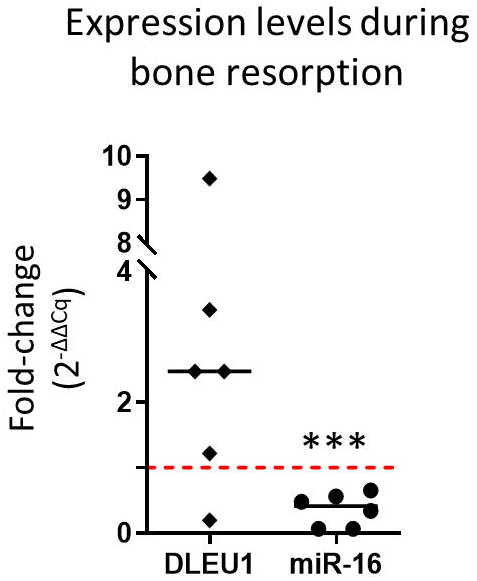
Expression levels of *DLEU1* and miR-16 during bone resorption. Mature OC were seeded onto bone slices at day 9 and left to resorb until day 12 (*N*=6).

**Supplemental Table I.**
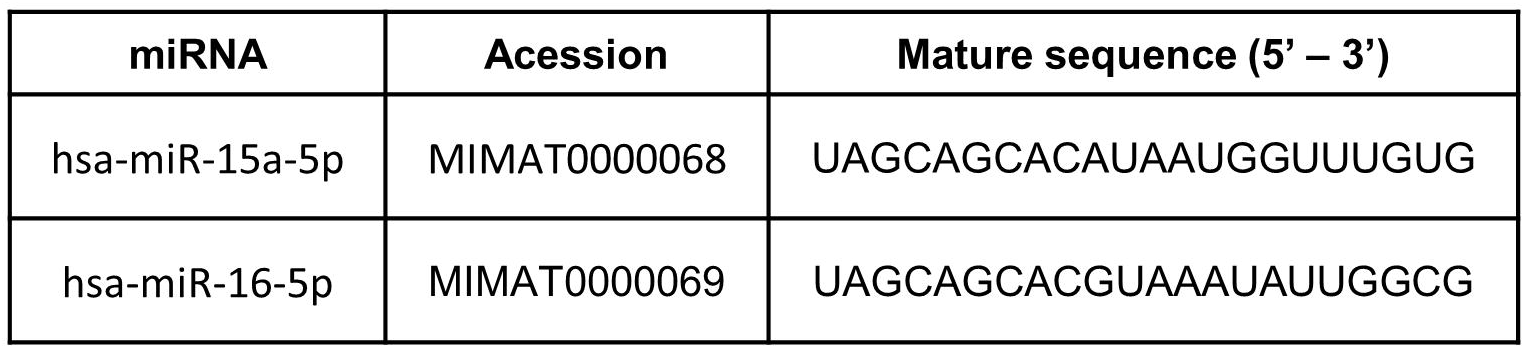
Mature miRNAs sequences according to miRbase annotations (http://www.mirbase.org/).

**Supplemental Table II.**
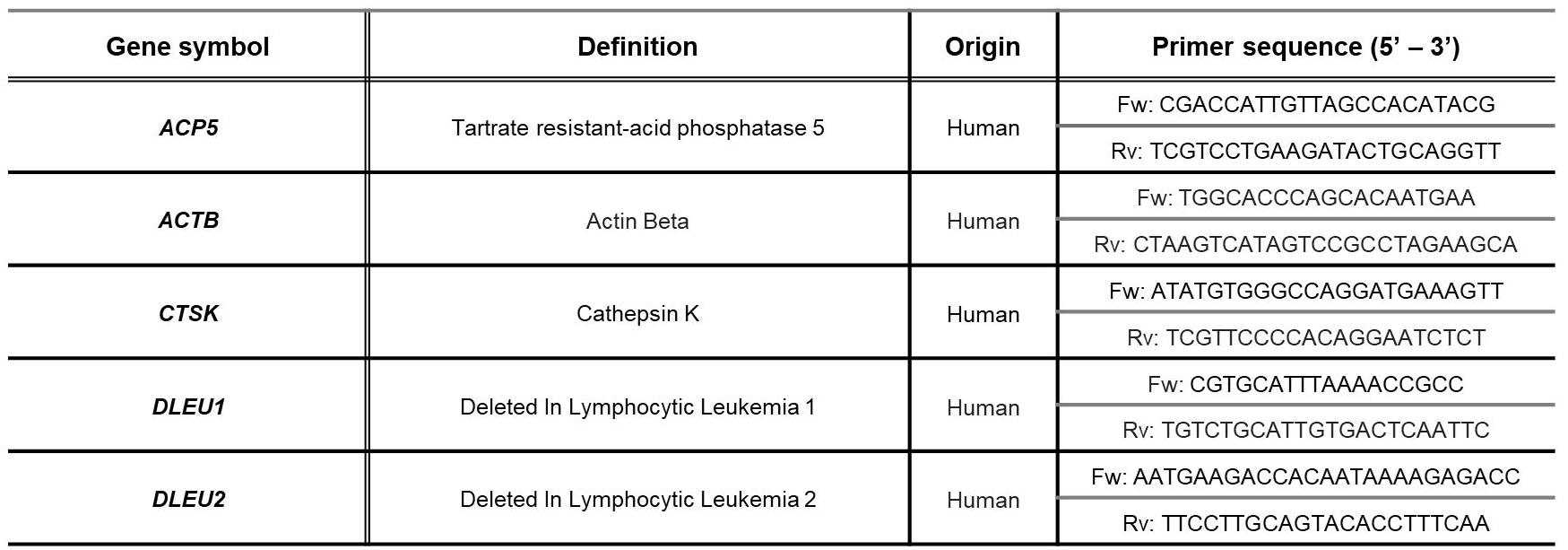
Primers used for reverse transcription quantitative real-time PCR.

